# Expanding GABAergic Neuronal Diversity in PSC-Derived Disease Models

**DOI:** 10.1101/2024.12.03.626438

**Authors:** Ruiqi Hu, Linda L. Boshans, Yiran Tao, Bohan Zhu, Junglong Li, Xindi Li, Rui Dang, Marianna Liang, Peiwen Cai, Mark Youssef, Gizem Inak, Yingnan Song, Xuran Wang, Alexander M Tsankov, Joseph D. Buxbaum, Peng Jiang, Zhiping P. Pang, Sai Ma, Nan Yang

**Author notes:** Correspondence (S.M.); (N.Y.). These authors contributed equally (Ruiqi Hu, Linda L. Boshans).

## Abstract

GABAergic interneurons comprise diverse molecular and functional subtypes that contribute to neural circuit assembly and are implicated in a range of neurodevelopmental and neuropsychiatric disorders. Traditional approaches for differentiating human pluripotent stem cells (PSCs) into neurons often face challenges such as incomplete neural differentiation, prolonged culture periods, and variability across PSC lines, which can constrain their application in disease modeling. To address these limitations, we developed a strategy that combines inducible expression of the transcription factors (TFs) ASCL1 and DLX2 with dual-SMAD and WNT inhibition to efficiently drive differentiation of human PSCs into diverse, region-specific GABAergic neuronal types. Using single-cell sequencing, we characterized the cellular heterogeneity of GABAergic induced neurons (iNs) generated with the patterning factors (patterned iNs) and those derived solely with TFs (PSC-derived iNs), revealing distinct interneuron subtype compositions and the regulatory programs that govern their fate specification. Patterned iNs exhibited gene expression features corresponding to multiple brain regions, particularly the ganglionic eminence (GE) and neocortex, whereas GABAergic PSC-derived iNs predominantly resembled hypothalamic and thalamic neurons. Both iN types were enriched for genes relevant to neurodevelopmental and psychiatric disorders, with patterned iNs more specifically linked to neural lineage genes, highlighting their utility for disease modeling. We further applied this protocol to investigate the impact of a recurrent ADNP syndrome-associated mutation (p.Tyr719*) on GABAergic neuron differentiation, revealing that this mutation disrupts GABAergic fate specification and synaptic transmission. Overall, this study expands the toolkit for disease modeling by demonstrating the complementary advantages of GABAergic PSC-derived iNs and patterned iNs in representing distinct GABAergic neuron subtypes, brain regions, and disease contexts. Together, these approaches provide a flexible platform for investigating molecular and cellular mechanisms relevant to neurodevelopmental and neuropsychiatric disorders.

## INTRODUCTION

Inhibitory GABAergic (γ-aminobutyric acid, GABA) neurons comprise a highly diverse class of cells defined by distinct molecular identities and are essential for the organization of neural circuits.^1,2^ Forebrain interneurons are generated in transient progenitor domains in the ventral telencephalon, known as the ganglionic eminences (GE), including the medial, caudal, and lateral ganglionic eminences (MGE, CGE, and LGE). These progenitor domains generate interneuron populations with diverse molecular and functional properties.^3,4^ MGE-derived interneurons predominantly mature into parvalbumin-expressing (PV; PVALB) or somatostatin-expressing (SST/SOM) cortical interneurons,^5–9^ whereas CGE generates vasoactive intestinal peptide (VIP)-expressing interneurons and populations expressing calretinin (CR; CALB1), LAMP5, and CCK.^6,10–12^ More recent studies indicate that dorsal LGE-derived interneurons also contribute to cortical inhibitory populations, particularly in frontal regions and deep white matter, and are marked by MEIS2, FOXP2, and PAX6,^13,14^ highlighting the multiple developmental origins underlying cortical interneuron diversity.

Perturbations in interneuron specification and function have been implicated in autism spectrum disorder (ASD), epilepsy, schizophrenia, Alzheimer’s disease (AD), and numerous other neurodevelopmental and psychiatric disorders.^3,4,15–17^ Importantly, interneurons arising from distinct developmental domains all appear to contribute to disease-relevant phenotypes, rather than any single lineage being uniquely engaged. Dysfunction of PV⁺ and SST⁺ interneurons, primarily derived from the MGE, has been linked to multiple neurological and neuropsychiatric disorders, including AD, ASD, schizophrenia, and epilepsy, and is often associated with disrupted inhibition, altered excitability, and imbalance in excitation/inhibition (E/I balance). Meanwhile, CGE-derived interneurons, which constitute a substantial fraction of cortical interneurons and are proportionally expanded in humans and non-human primates^18–22^, have been implicated across multiple neuropsychiatric conditions^23–25^, and the CGE is the primary source of tumors and lesions in tuberous sclerosis complex (TSC)^26^. Together, these findings underscore the importance of modeling GABAergic neuron diversity and of developing in vitro systems capable of generating interneurons representing a broad range of subtypes and developmental origins.

Human pluripotent stem cell (PSC)-based approaches have enabled the scalable generation of GABAergic neurons and are widely used for mechanistic and disease-focused studies. Early conventional differentiation strategies largely relied on small-molecule and morphogen-based patterning to mimic MGE development, reflecting foundational insights from rodent studies in which MGE-derived interneurons constitute major cortical inhibitory populations. ^27^ Despite their utility, these approaches face several limitations, including partial induction of neural identities, protracted timelines required to generate mature neuronal cultures, limited representation of CGE- and LGE-derived subtypes, and variability in differentiation efficacy across PSC lines. In contrast to conventional paradigms, direct reprogramming of PSCs into defined cell types through forced expression of transcription factors (TFs) offers a compelling alternative. This method has been widely adopted because of its simplicity, speed, efficiency, scalability, and reproducibility. For instance, forced expression of proneural TFs such as Neurogenin-2 (NEUROG2 or NGN2) and achaete-scute complex-like 1 (ASCL1) - key regulators of neuronal fate during brain development - can rapidly induce neuronal differentiation in human PSCs. ^27–31^ While effective at generating neurons, TF induction alone is limited in its ability to produce the full spectrum of neuronal subtypes in the brain. Recent studies have addressed this limitation by combining TF overexpression with extrinsic patterning signals, enabling the specification of a broader range of neuronal subtypes that cannot be generated using either method alone. Notably, pre-patterning PSCs prior to NGN2 induction allows the generation of regionally specified neuronal populations, such as cortical-like or motor neurons.^32,33^ Despite this progress, comparable combinatorial strategies have not been systematically explored for generating diverse and developmentally relevant GABAergic interneuron subtypes.

Previously, we demonstrated that forced expression of Ascl1 and Dlx2 efficiently induces the differentiation of human PSCs into GABAergic neurons. These GABAergic induced neurons (iNs) robustly adopt inhibitory neurotransmitter phenotypes and express subtype-specific markers, indicating the generation of potentially diverse subpopulations through this protocol.^27^ We and others have shown that GABAergic iNs are valuable for investigating inhibitory synapse-specific phenotypes linked to human neurological disorders.^27,34,35^ However, the characterization of GABAergic iNs has predominantly relied on electrophysiology, limited immunofluorescence markers, and bulk transcriptomic profiling. As a result, their precise subtype composition, regional correspondence to *in vivo* interneuron populations, and degree of heterogeneity remain poorly defined at single-cell resolution.

To address these outstanding issues, we demonstrate in this study that neuralization of stem cells via dual SMAD and WNT inhibition^36^, followed by forced expression of ASCL1 and DLX2, effectively specifies diverse GABAergic neuronal identities that correspond to populations found in the developing forebrain. Using single-cell RNA sequencing (scRNA-seq), we comprehensively characterize GABAergic iNs generated from small-molecule–patterned PSCs (patterned-iNs) and compare them with PSC-derived ASCL1/DLX2-iNs (PSC-iNs) and GABAergic neurons generated using small-molecule patterning alone. This comparative analysis reveals that each differentiation strategy yields a diverse array of GABAergic neuronal subtypes with composite regional identities. We also investigate the differentiation dynamics and regulatory mechanisms underlying fate specification in PSC-iNs and patterned-iNs using single-cell multiomic approaches. To further demonstrate the utility of patterned GABAergic iNs, we apply this framework to *ADNP* (activity-dependent neuroprotective protein), one of the most frequently mutated genes in syndromic autism.^37^ The recurrent truncating mutation p.Tyr719* is the most common pathogenic ADNP variant identified in patients. While prior studies of ADNP have mainly focused on excitatory neuron models, the effects of ADNP mutations in human inhibitory neurons, a lineage central to E/I balance, remain poorly characterized. Using patterned GABAergic iNs, we examine how the ADNP p.Tyr719* mutation influences interneuron subtype specification and inhibitory synaptic function. Together, this work demonstrates the complementary strengths of PSC-derived and patterned GABAergic neuron models in capturing interneuron diversity across brain regions and disease contexts, providing a versatile platform for investigating molecular and cellular mechanisms underlying neurodevelopmental and neuropsychiatric disorders.

## RESULTS

### ASCL1 & DLX2 Induce GABAergic Neurons from Patterned Progenitors

Direct programming leverages one or more TFs to force the establishment of a distinct transcriptional network, thereby promoting the adoption of a new cellular identity.^38,39^ Previously, we reported that forced expression of ASCL1 and DLX2 in human PSCs induces rapid differentiation into GABAergic neurons (Figure 1A). This strategy has since been adopted and independently validated by multiple groups to generate human GABAergic neurons.^40,41^ While these iN cells displayed core features of GABAergic neurons, including GAD1/2, DLXs, and SLC32A1 (vGAT), the homogeneity of the resulting cell population and its similarity to *in vivo* counterparts remained unclear. To address these questions, we performed single-cell RNA sequencing (scRNA-seq) to profile GABAergic PSC-iNs 42 days after ASCL1 and DLX2 expression (Figure 1A).

**Figure 1.**
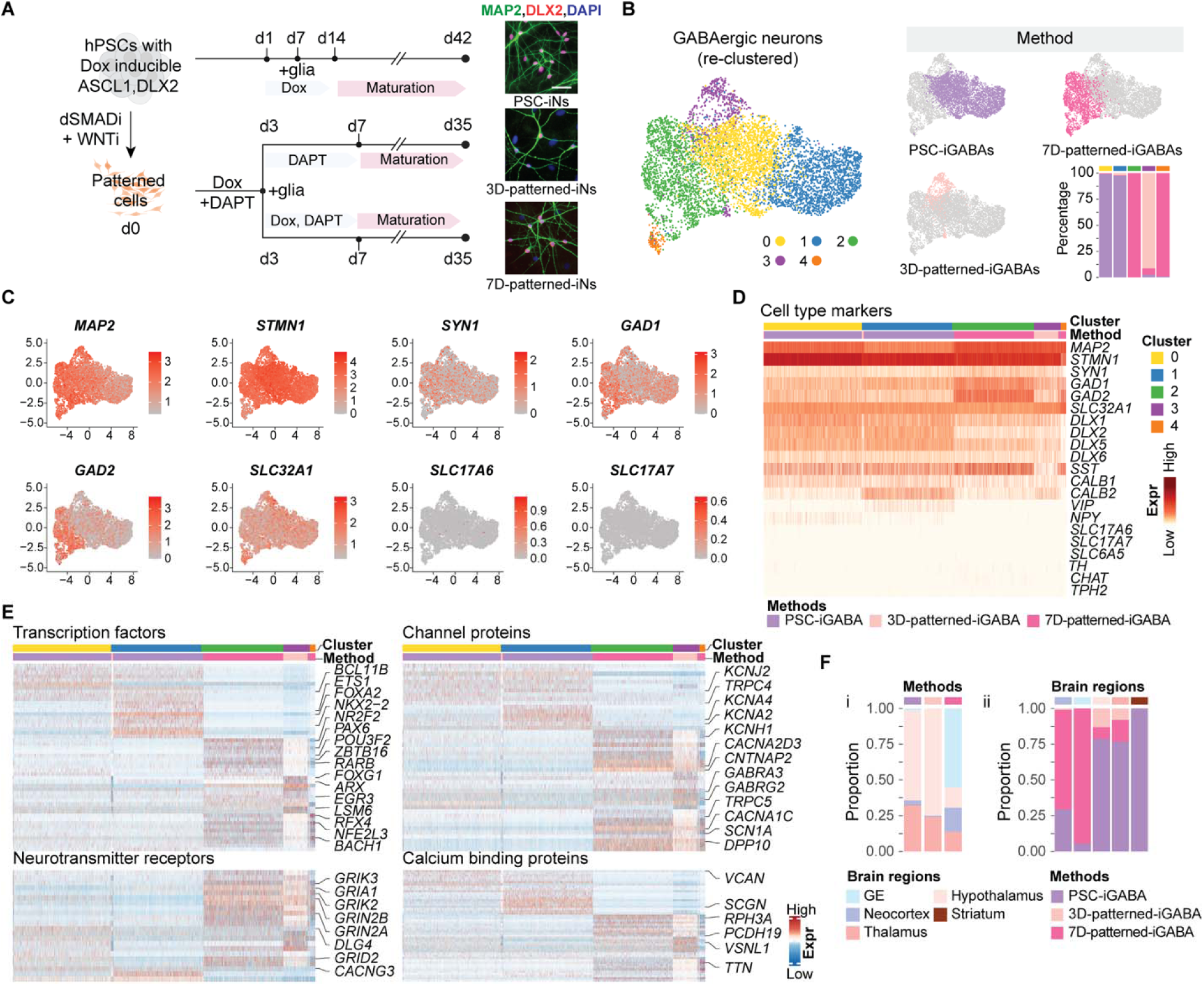
Generation of Diverse GABAergic iNs Using Different Strategies. (**A**) Schematic of the differentiation protocols used to generate GABAergic iNs from hPSCs using either TFs only (PSC-iGABA) or small molecule patterning followed by ASCL1/DLX2 induction (3D- and 7D-patterned-iGABAs). Representative immunofluorescence images show co-expression of MAP2 and DLX2 in all three conditions. Scale bar, 50 µm. Abbreviations: dSMADi (dual SMAD inhibition), WNTi (WNT inhibition), PSC-iNs (PSC-derived induced neurons), 3D-patterned-iNs (patterned induced neurons with 3 days of doxycycline exposure), and 7D-patterned-iNs (patterned induced neurons with 7 days of doxycycline exposure). (**B**) UMAP embedding of *SCL32A1*^+^ GABAergic iNs, color-coded by cluster identity and differentiation method. Right: Cluster distribution for each method, with corresponding bar plot showing the percentage of cells from each condition in each cluster. (**C**) Feature plots showing inhibitory and excitatory neuron marker gene expression in GABAergic-induced neurons. (**D, E**) Heatmap of cell type-specific markers, TFs, channel proteins, neurotransmitter receptors, and calcium-binding protein expression within each GABAergic iN cluster. The selected genes are implicated in psychiatric and neurological conditions, with a particular focus on ASD and epilepsy. (**F**) i. Proportions of cells from each method mapped to different brain regions using label transfer from primary human fetal brain datasets. ii. Contribution of each method to cells annotated as each brain region. Aggregated proportions showing that GE- and cortex-like cells are most enriched in 7D-patterned-iNs, whereas PSC-iGABAs show higher mapping to the hypothalamus and thalamus. See also Figures S1, S2, S3, Table S2.

For comparison, we generated interneurons using a well-established directed differentiation protocol that involves dual SMAD inhibition, WNT signaling inhibition, and SHH signaling activation to specify ventral telencephalic progenitors that give rise to interneurons.^42–45^ This approach is presumed to produce GABAergic neurons similar to PSC-iNs, as evidenced by the expression of interneuron subtype markers such as SST, CR, and Calbindin (CB). ^27,42^ We characterized these neurons five weeks after culturing the progenitors on dissociated mouse glial cells (Figure S1A). While neurons generated using the directed differentiation method expressed GABAergic markers like *GAD1* and *DLX6-AS*, they showed minimal expression of markers associated with synaptic function, including *SLC32A1* and *SYN1*. In contrast to PSC-iNs, these neurons expressed *FOXG1*, a defining marker of forebrain identity, suggesting that they represent immature cortical interneurons (Figures S1B and S1C). The lack of FOXG1 expression in PSC-iNs indicates that they may resemble GABAergic neurons from more posterior brain regions.

These observations prompted us to test whether extrinsic patterning cues could synergize with ASCL1 and DLX2 to rapidly generate diverse inhibitory neurons associated with specific brain regions. To this end, we employed a doxycycline (DOX)-responsive piggyBac transposon system to achieve robust and controlled overexpression of ASCL1 and DLX2 in human cells. We first induced central nervous system neuroectoderm from human PSCs following an established forebrain patterning timeline, inhibiting TGF-β and BMP signaling with SB431542 and LDN193189 for 10 days, while transiently blocking WNT signaling with the tankyrase inhibitor XAV939 during the first 3 days.^46^ This initial patterning phase biases cells toward a forebrain neural fate but does not by itself specify interneuron identity. Following this patterning phase, DOX was applied to induce ASCL1 and DLX2 expression, together with the γ-secretase inhibitor DAPT, to eliminate dividing progenitors and synchronize maturation.^47^ Cells were then dissociated and co-cultured with mouse astrocytes to support neuronal maturation and synapse formation. To assess whether the duration of ASCL1 and DLX2 expression influences neuronal fate, DOX was withdrawn after either 3 or 7 days post-induction, generating 3D-patterned-iNs and 7D-patterned-iNs, respectively. We refer to these cells as patterned GABAergic induced neurons (patterned-iNs) to reflect the combination of brief regional patterning and transcription factor–driven fate specification. Five weeks after astrocyte co-culture, GABAergic neurons expressing MAP2 and DLX2 were detected under all experimental conditions (Figure 1A).

To gain deeper insights into the cellular composition of these cultures, we performed scRNA-seq on neurons generated under each differentiation condition and analyzed them alongside GABAergic PSC-iNs. In total, we generated transcriptomes from 12,876 cells across various differentiation strategies, including 5,314 PSC-iNs, 2,899 3D-patterned-iNs, and 4,663 7D-patterned-iNs that passed quality control (Table S1). Dimensionality reduction and visualization using uniform manifold approximation and projection (UMAP) revealed seven molecularly distinct cell clusters that segregated by differentiation strategy (Figure S2A). Further examination of neuronal marker gene expression revealed that most cells exhibit neuronal characteristics as marked by *MAP2, STMN1,* and *SYN1* expression (Figures S2B and S2C). Most of the induced neurons (86.2%) are GABAergic, as indicated by the expression of *GAD1*, *GAD2*, *SLC32A1*, and *DLXs*. A small fraction of cells was identified as glutamatergic, based on *SLC17A6/7* expression (3D-patterned-iNs: 42/2,899; 7D-patterned-iNs: 56/4,663; PSC-iNs: 10/5,314), while only rare cells expressed markers of other neuronal lineages, including CHAT (cholinergic, 0.02%) or TH (dopaminergic, 0.06%). Notably, the proportion of cells expressing the vesicular GABA transporter *SLC32A1* was substantially lower in 3D-patterned-iNs compared with 7D-patterned-iNs (19.4% versus 50.9%). Conversely, 3D-patterned-iNs showed increased expression of genes associated with a less mature or progenitor-like state, including *SOX2* (7.0% versus 0.94% in 7D-patterned-iNs). These data suggest that 3 days of ASCL1 and DLX2 expression are insufficient to fully drive small-molecule patterned progenitor-like cells into a mature neuronal state. Together, our data demonstrate that the duration of TF expression, in combination with extrinsic patterning cues, critically influences neuronal maturation and supports the synergistic use of TFs and small molecules to generate GABAergic neurons from human PSCs.

### Molecularly Diverse GABAergic Neurons Generated by Distinct Differentiation Strategies

To further investigate heterogeneity among induced GABAergic neurons (iGABAs), we extracted *SLC32A1*-expressing neurons, comprising 4,086 PSC-iGABAs, 574 3D-patterned-iGABAs, and 1,915 7D-patterned-iGABAs. Unsupervised clustering based on gene expression identified five major molecular clusters, with clear separations between clusters depending on the differentiation method used (Figure 1B). As expected, all clusters consistently expressed canonical markers of the GABAergic lineage while generally devoid of markers indicative of other neuronal lineages (Figures 1C and S3A). Consistent with our previously reported immunofluorescence analysis, these clusters showed variable enrichment for inhibitory neuron subtype markers, including *SST*, *CALB1*, *CALB2*, *VIP,* and *NPY* (Figures 1C, 1D, and S3A).^27^ When examining the differentially expressed TFs across these clusters, we found *FOXG1* expressed in patterned-iGABAs (clusters 3 and 4). The gene expression signature of 7D-patterned-iGABAs (cluster 2) reflected spatial origins and subtypes associated with the CGE and LGE, marked by *NR2F2* (*COUP-TFII*), *PAX6*, and *ISL1*.^13,48,49^ *ISL1* was also strongly expressed in cluster 0, which consisted of PSC-iGABAs, whereas PSC-iGABAs in cluster 1 robustly expressed *FOXA2* and *NKX2-2*, markers associated with hypothalamic inhibitory neurons.^50,51^ In addition to TFs, we identified differentially expressed genes (DEGs) encoding ion channels, neurotransmitter receptors, and calcium-binding proteins across molecularly distinct clusters, further underscoring the molecular diversity generated by distinct differentiation strategies (Figure 1E and Table S2). Notably, expression of the NMDAR subunits *GRIN2A* and *GRIN2B* was higher in neurons produced with the small-molecule patterning (Figure S3B). *GRIN2B* is predominantly expressed during prenatal cortical development, whereas *GRIN2A* expression emerges largely postnatally.^52^ This pattern aligns with prior reports demonstrating that combining regional patterning with lineage-specific TF induction promotes NMDAR subunit expression^33^, suggesting a synergistic effect on neuronal maturation.

To annotate the brain regional identities represented by the iGABAs, we applied a reference-mapping approach using scRNA-seq datasets from primary human fetal brain regions,^53,54^ including neocortex, GE, striatum, thalamus, hypothalamus, midbrain, and claustrum (see Methods). iGABAs mapped across multiple forebrain regions, including the hypothalamus, thalamus, neocortex, and GE (Figure 1F). While all differentiation strategies produced neurons spanning multiple regional identities, GE- and neocortex-associated populations were enriched in patterned ASCL1/DLX2 conditions (7D-patterned-iGABAs), whereas PSC-iGABAs generated without small-molecule patterning preferentially aligned with hypothalamic and thalamic identities. These assignments were supported by regional marker expression, including *FOXA2* in hypothalamic clusters^50,55^ and *NR2F2* in the GE-associated cluster (Figure S3C).

Next, to benchmark interneuron diversity across differentiation strategies, we performed comparative single-cell transcriptomic analyses including 7D-patterned-iGABAs, PSC-iGABAs, and GABAergic neurons generated using previously published small-molecule–based protocols reported by other groups^45,56^. All datasets were uniformly annotated using the Feng et al. (2025) human developmental brain atlas, which resolves interneuron subtypes derived from MGE, LGE, and CGE origins across cortical and subcortical regions.^49^ Across methods, human PSC-derived GABAergic neurons mapped to MGE-, CGE-, and LGE-associated lineages (Figure S4). However, neurons generated using small-molecule patterning protocols were strongly skewed toward MGE-derived interneuron subtypes, with limited representation of CGE- and LGE-associated populations. In contrast, both PSC-iGABAs and patterned-iGABAs spanned multiple developmental origins, yet interneurons assigned to the same MGE, CGE, or LGE domains occupied distinct subtype states depending on the differentiation strategy, indicating that protocol choice shapes fine-grained interneuron identity beyond broad lineage assignment.

### Functional Maturation and Circuit Integration of Patterned GABAergic iNs

Given transcriptomic evidence that 3D-patterned GABAergic iNs retain immature features, we focused subsequent analyses on 7D-patterned GABAergic induced neurons (hereafter referred to as GABAergic patterned iNs or patterned-iGABAs). To assess functional maturation *in vitro*, we performed whole-cell patch clamp recordings and immunofluorescence analyses at day 35 of differentiation (Figure 2A). Voltage-clamp recordings revealed prominent inward sodium currents in response to depolarizing steps (Figure 2B), and current-clamp traces showed robust trains of action potentials with increasing current injection (Figure 2C). Patterned-iNs exhibited passive membrane properties consistent with neuronal maturity (Figure 2D). To assess functional inhibitory synapse formation, we recorded spontaneous inhibitory postsynaptic currents (sIPSCs). sIPSCs were recorded in the presence of CNQX and abolished by application of picrotoxin, confirming GABAergic synaptic transmission (Figures 2E and 2F). Immunostaining further confirmed interneuron subtype diversity, including SST⁺, CR⁺, and CB⁺ populations, together with expression of the cortical identity marker FOXG1 and the CGE-associated marker PAX6 (Figure 2G). Together, these results indicate that patterned-iN cultures generate functionally mature GABAergic neurons with diverse molecular, regional, and subtype identities.

**Figure 2.**
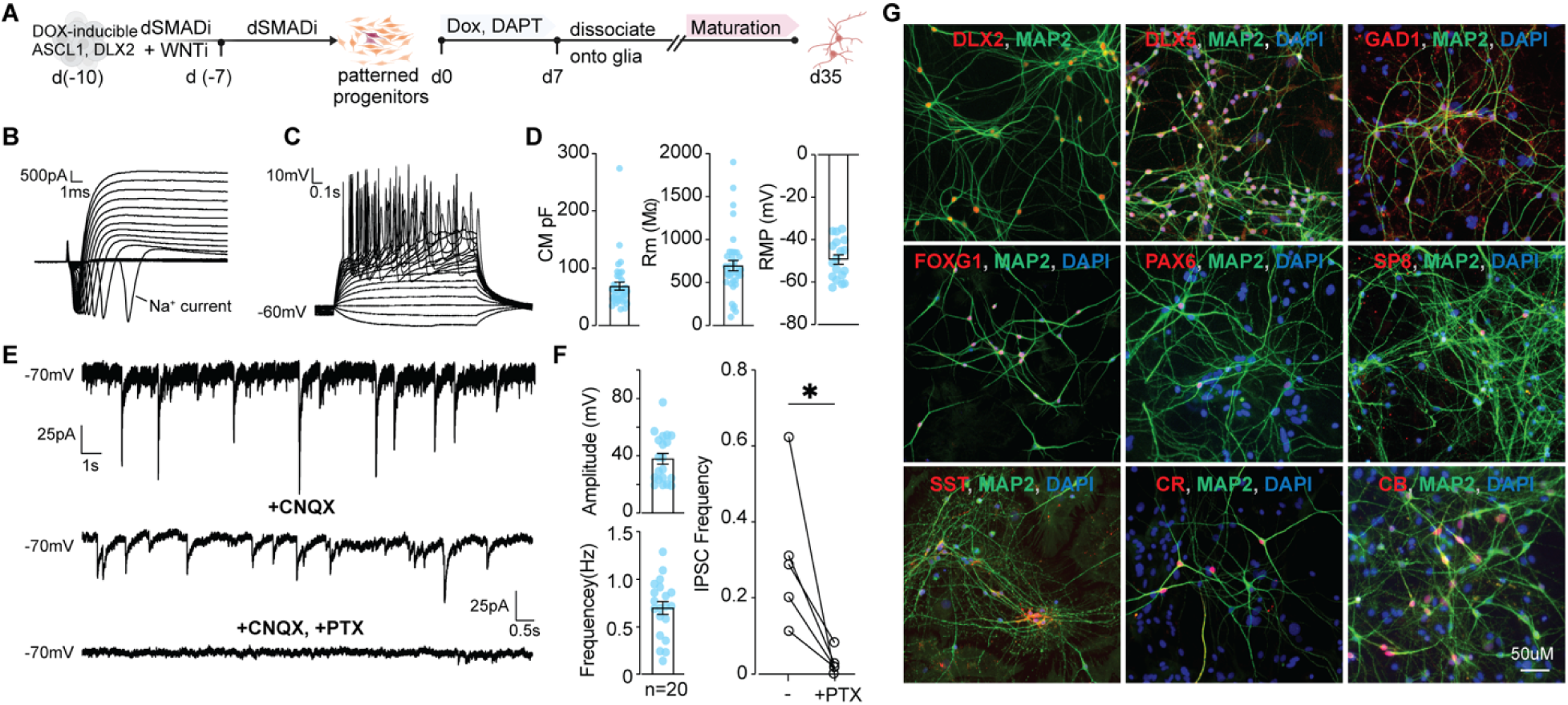
Functional and Subtype Characterization of patterned-iGABA Neurons. (**A**) Schematic of the differentiation protocol. (**B**) Voltage-clamp recording of inward sodium currents evoked by step depolarizations. (**C**) Current-clamp recording showing repetitive action potential firing across injected current steps. (**D**) Summary of passive membrane properties: membrane capacitance (CM) (n = 38), input resistance (Rm) (n = 38), and resting membrane potential (RMP) (n = 20). Each dot represents an individual recorded neuron. (**E**) Voltage-clamp recordings of spontaneous inhibitory postsynaptic currents (sIPSCs) at −70LmV. CNQX partially blocks EPSCs, and complete blockade is achieved with CNQX + picrotoxin (PTX). (**F**) Quantification of sIPSC amplitude and frequency across individual neurons (n = 20), with a significant decrease in IPSC frequency following PTX treatment (p < 0.05). (**G**) Immunofluorescence staining confirms the presence of multiple interneuron subtypes, including GAD1+, PAX6+, FOXG1+, SST+, CR+, and CB+ cells, co-labeled with MAP2. Scale bar: 50Lµm.

To determine whether patterned-iNs can survive, mature, and integrate into neural circuits *in vivo*, we transplanted EGFP-labeled immature patterned-iNs into the periventricular region of neonatal Rag2⁻/⁻ mice. At five months post-transplantation, immunohistological analysis revealed that the majority of human nuclear antigen (HNA)–positive cells remained near the injection site and differentiated into NeuN⁺ neurons, indicating stable neuronal maturation *in vivo* (Figures 3A and S5). To assess functional integration into host circuits, we performed whole-cell patch-clamp recordings from GFP⁺ human neurons in acute cortical slices. Across 12 recorded cells from three animals, depolarizing current injections elicited robust action potential firing, with a mean rheobase of 30 ± 14 pA and firing frequencies reaching 42 ± 8 Hz at higher current steps (Figures 3B-D, Table S3). Action potentials exhibited mature biophysical properties, and passive membrane properties were consistent with mature functional interneurons (Figures 3E–J). Importantly, the human neurons received spontaneous synaptic inputs, including both excitatory postsynaptic currents (EPSCs) and inhibitory postsynaptic currents (IPSCs) (Figures 3K and 3L, Table S3), demonstrating synaptic connectivity with host cortical circuits.

**Figure 3.**
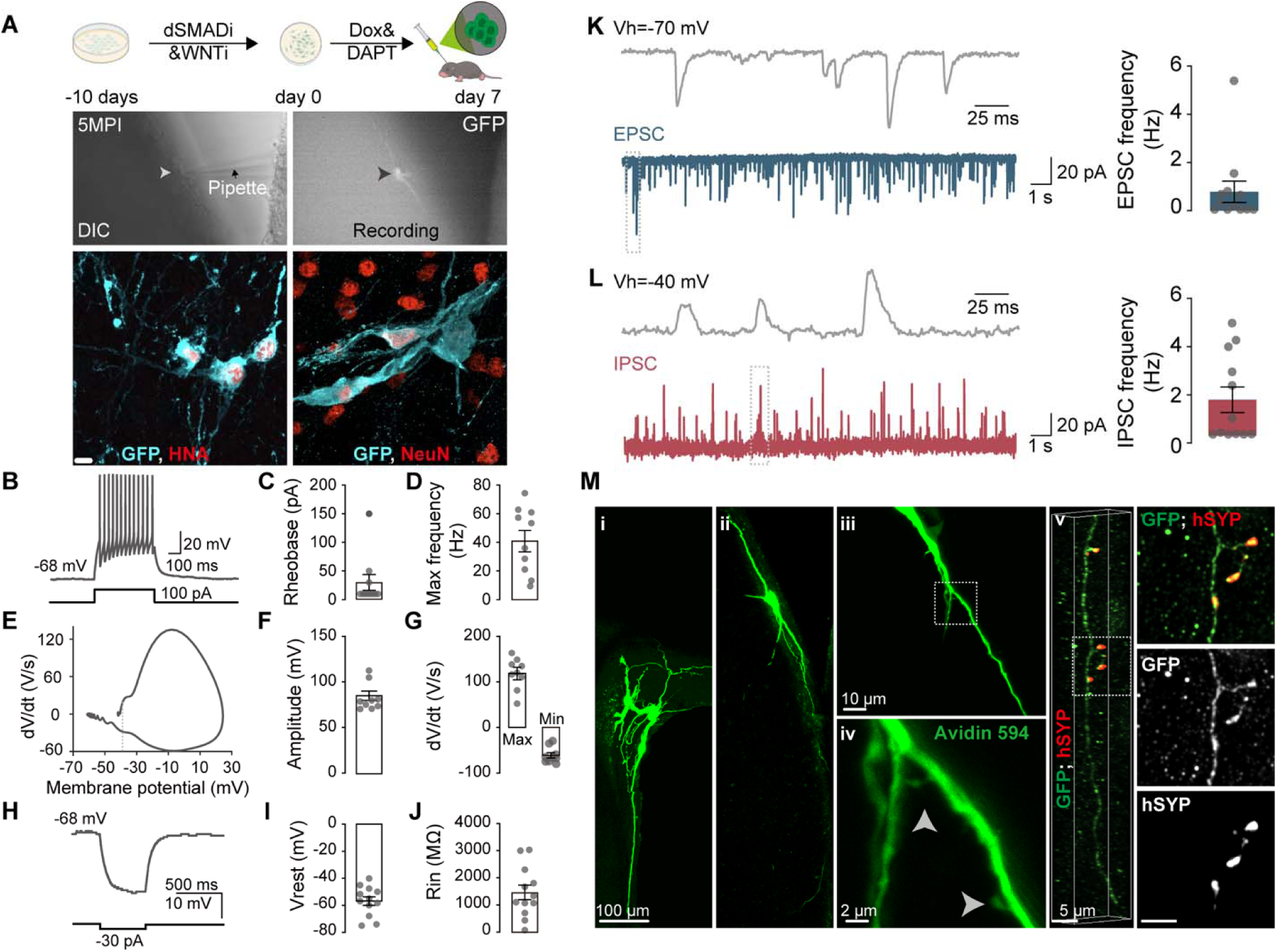
Functional and morphological characterization of human iPSC-derived interneurons following transplantation into the mouse cortex. (**A**) Top: Schematic overview and experimental timeline. Human iPSCs were patterned with dual SMAD and WNT inhibition (dSMADi & WNTi) for 10 days, followed by doxycycline (Dox) and DAPT treatment for 7 days, and subsequently transplanted into the cortex of postnatal day 2 (P2) immunodeficient mice. Whole-cell patch-clamp recordings were performed approximately 5 months post-injection (MPI). Middle left: Differential interference contrast (DIC) image of an acute brain slice showing a patch pipette approaching a GFP+ human induced neuron (iN). Arrowhead indicates the soma targeted for recording. Middle right: Fluorescent image of the same GFP+ neuron during patch-clamp recording. Bottom: Confocal images showing the GFP+ neurons (cyan) co-labeled with human nuclear antigen (HNA) or NeuN (red), confirming human neuronal identity. Scale bar: 5 μm. (**B**) Representative current-clamp recording from a GFP+ human neuron showing APs in response to a 500 ms, 100 pA current injection. Scale: 20 mV, 100 ms. (**C**) Group data showing rheobase values from 10 neurons (3 mice). (**D**) Maximum firing frequency of APs across the same group of GFP+ neurons. (**E**) Example phase-plane plot (dV/dt vs. membrane potential) of a single AP from a GFP+ cell. (**F**) Group analysis of AP amplitude (left) and the maximal and minimal dV/dt slopes (right). (**G**) Voltage response of a GFP+ neuron to a 500 ms, −30 pA hyperpolarizing current step. Scale: 10 mV, 500 ms. (**H**) Group data showing resting membrane potential (V_rest) and input resistance (R_in) of GFP+ neurons (n=12 cells from 3 mice). (**K, L**) Representative voltage-clamp recordings showing spontaneous excitatory (**K**) and inhibitory (**L**) postsynaptic currents in GFP+ human neurons. Excitatory postsynaptic currents (EPSCs, blue) were recorded at a holding potential of –70 mV, while inhibitory postsynaptic currents (IPSCs, red) were recorded at a holding potential of –40 mV. Insets display expanded views of boxed regions to highlight individual synaptic events. Scale bars: 20 pA, 1 s (main traces), 25 ms (insets). Right panels: Quantification of spontaneous EPSC and IPSC frequencies in GFP+ neurons (n = 12 cells each). (**M**) Morphological reconstruction and synaptic marker localization in transplanted human interneurons. (**i–iv**) Biocytin–Avidin 594 staining of whole-cell recorded human GFP+ neurons reveals detailed cellular morphology. (i-ii) Overview of multiple recorded human neurons in the transplant region, filled with biocytin and visualized with Avidin 594 (green). (iii) High-magnification image of a single biocytin-filled neuron, showing its soma and dendritic arbor. (iv) Further magnified view highlights fine dendritic structures and spine-like protrusions (arrowheads). (v) Confocal z-projection showing co-localization of human-specific synaptophysin (hSYP, red), a presynaptic marker, along dendrites of a GFP+ interneuron (green). Split channels are shown below to confirm punctate hSYP localization on GFP-labeled neuronal processes. Scale bars: 100 μm (**i**), 10 μm (**ii–iii**), 2 μm (**iv**), 5 μm (**v**). Data are represented as mean ± SEM. See also Figure S5 and Table S3.

**Figure 4.**
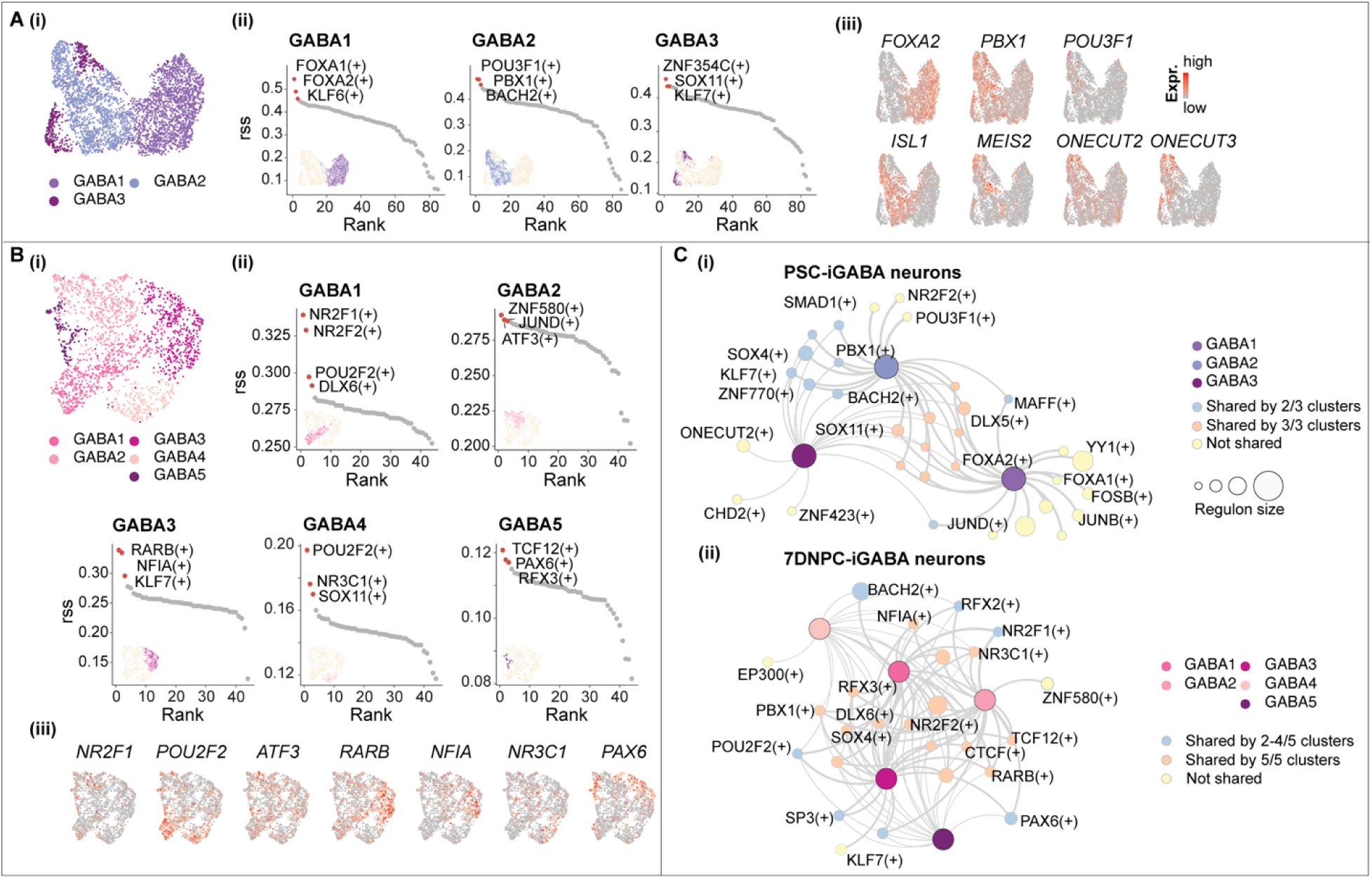
Gene Regulatory Networks Underlying Induced GABAergic Neuron Diversity. (A-B) Presented are the following for the GABAergic PSC-iNs (A) and the patterned iNs (B): (i) identified clusters, (ii) ranked regulons, and (iii) UMAP visualization of the top TFs associated with GABAergic neuron fate specification. **(C)** GRN analysis focused on cluster-specific modulators for **(i)** GABAergic PSC-iNs and **(ii)** patterned iNs. Regulons are color-coded by uniqueness, with node size proportional to the number of target genes regulated by each TF. Edge thickness indicates the strength of the association between a cluster modulator and its corresponding regulon. See also Figure S7.

Finally, immunohistochemical analysis using a human-specific synaptophysin antibody revealed presynaptic puncta associated with GFP⁺ human neurons, providing additional evidence of synaptic connectivity *in vivo* (Figure 3M). Transplanted cells also displayed complex neuronal morphologies with elaborated dendritic arbors. Together, these findings demonstrate that patterned-iNs not only acquire functional maturity in vitro but also survive long term, receive synaptic inputs, and integrate into host cortical circuits following transplantation.

### Neuropsychiatric and Neurological Disease Gene Enrichment Across GABAergic iN Types

Induced neurons offer significant potential for studying neuropsychiatric and neurological diseases *in vitro* and for linking genetic risk to cellular phenotypes, offering valuable insights for basic research and pharmaceutical development. To evaluate the disease relevance of different GABAergic iN populations, we performed gene set enrichment analysis using Disease Enrichment (DisGeNET)^57^ on shared and condition-specific genes expressed in PSC- or patterned-iGABAs. Our analysis revealed that genes shared between PSC- and patterned-iGABAs were strongly enriched for risk genes associated with various brain disorders, including intellectual disability, neurodevelopmental disorders (NDDs), schizophrenia, epilepsy, and bipolar disorder (Figure S6A; Table S4; q-value ≤ 0.05). Notably, genes specific to patterned iNs also showed strong enrichment in psychiatric and neurological conditions, whereas genes specific to PSC-iNs were preferentially enriched for non-neural disease annotations (Figures S6B and S6C). Consistent with this observation, patterned-iN-specific genes also exhibited robust enrichment for ASD risk genes curated in the SFARI database^58^ (odds ratio = 2.50; BH-corrected *p* = 1.05 × 10⁻¹³). These findings indicate that while both PSC- and patterned iNs are suitable for investigating disease-associated genotype-phenotype relationships, patterned iNs exhibit closer alignment with neural lineages, making them particularly advantageous for modeling neural-related diseases. To further extend this analysis, we incorporated comparative transcriptomic mapping using publicly available scRNA-seq datasets derived from postmortem brain tissue of individuals with AD and ASD.^59,60^ This approach enabled us to assess whether the transcriptional identities of induced GABAergic neurons correspond to interneuron populations implicated in these disease contexts. These analyses revealed overlap between patterned-iNs and disease-associated interneuron signatures observed in AD and ASD datasets, supporting their relevance for modeling disease-linked inhibitory neuron phenotypes (Figure S6D).

### Gene Regulatory Networks Underlying Diverse GABAergic-Induced Neurons

Next, to explore the regulatory mechanisms underlying GABAergic neuron identities generated by different differentiation strategies, we applied single-cell regulatory network inference and clustering (SCENIC), which infers TF–centered regulons and their target gene networks from scRNA-seq data.^61^ This analysis identified both shared and strategy-specific regulons across PSC-iGABAs and patterned-iGABAs, revealing distinct gene regulatory architectures associated with each differentiation approach (Figure S7). At a global level, regulons associated with PSC-iGABAs included well-recognized determinants of interneuron cell fate and migration, such as DLX2 and DLX5. Regulons linked to 7D-patterned-iGABAs, such as NR2F1 (COUP-TFI), NR2F2 (COUP-TFII), and PAX6, are recognized for their roles in the differentiation of specific interneuron subtypes, including those originating from the LGE and CGE ^13,62,63^.

Because relatively few GABAergic neurons were recovered following 3-day DOX induction, we focused subsequent analyses on PSC-iNs and 7D-patterned iNs to identify specifically activated regulons within each molecular cluster. Neurons from these conditions were subdivided into molecular clusters based on transcriptomic profiles, and regulons were ranked using the Regulon Specificity Score (RSS) to identify cluster-enriched regulatory programs (Figures 3A and 3B). Our analysis revealed that each neuronal cluster was associated with different sets of regulons. Specifically, within the GABAergic PSC-iNs, prominent regulons included FOXA1, FOXA2, and PBX1, TFs associated with hypothalamic regionalization, neurogenesis, and neuroendocrine function. FOXA1 and FOXA2 emerged as primary regulons in cluster 1, consistent with their established expression in the embryonic hypothalamus and its role in regulating adaptive responses to metabolic state, such as fasting.^50,55^ PBX1 has similarly been implicated in hypothalamic molecular pathways involved in hormonal regulation.^64^ We also showed the expression patterns of these top-ranked regulators in corresponding neuronal clusters (Figure 3A). In addition, consistent with our observations that subsets of PSC-iNs exhibit striatal-like features, several highly expressed TFs associated with both striatal and hypothalamic differentiation and function were identified, including ONECUTs, MEIS2, and _ISL1.48,65,66_

In contrast, we found that subclusters of GABAergic neurons generated with patterning factors were enriched in a distinct set of regulons associated with the processes of neural fate determination and migration, including *NR2F1/2, DLX6*, *RARB*, *POU2F2*, *TCF12,* and *PAX6* (Figure 3B). Notably, NR2F1, NR2F2, and PAX6 are expressed in GE and play key roles in the development of cortical and ventromedial forebrain inhibitory neurons. Compared with PSC-iNs, patterned-iNs exhibited more homogeneous regulatory programs across clusters, reflected by broader sharing of key regulons and reduced cluster-specific segregation (Figure 3C). Together, these analyses demonstrate that differentiation strategy shapes GABAergic neuron identity not only at the level of transcriptomic subtype composition but also through distinct gene regulatory networks, highlighting how TF–based induction and extrinsic patterning cues engage overlapping yet non-identical regulatory programs to generate diverse inhibitory neuron states.

### Distinct Lineage-Specific Gene Regulatory Networks Induced by ASCL1&DLX2 in PSCs and Patterned Progenitors

Epigenetic regulation plays a pivotal role in determining and specifying cell lineages. To gain insights into the regulatory mechanisms that drive the fate specification of PSC-iNs and patterned-iNs, we performed SHARE-seq (Simultaneous High-throughput ATAC and RNA Expression with Sequencing)^67^ on PSCs, patterned progenitors, and cells at the early stages following ASCL1 and DLX2 induction (Figure 5A). This approach enabled simultaneous profiling of chromatin accessibility and gene expression within individual cells, providing a comprehensive view of the regulatory landscape underlying neuronal differentiation. We obtained high-quality ATAC-seq and RNA-seq data from 23,434 cells. To reveal global similarities and distinctions among individual cells, we applied dimensionality reduction using Latent Semantic Indexing (LSI) to single-cell ATAC-seq data and PCA to scRNA-seq data, followed by visualization using UMAP embedding. Chromatin accessibility and transcriptional representations were well aligned, revealing coherent trajectories that reflect the progression of cell states over time and segregate according to cell type of origin (Figure 5B).

**Figure 5.**
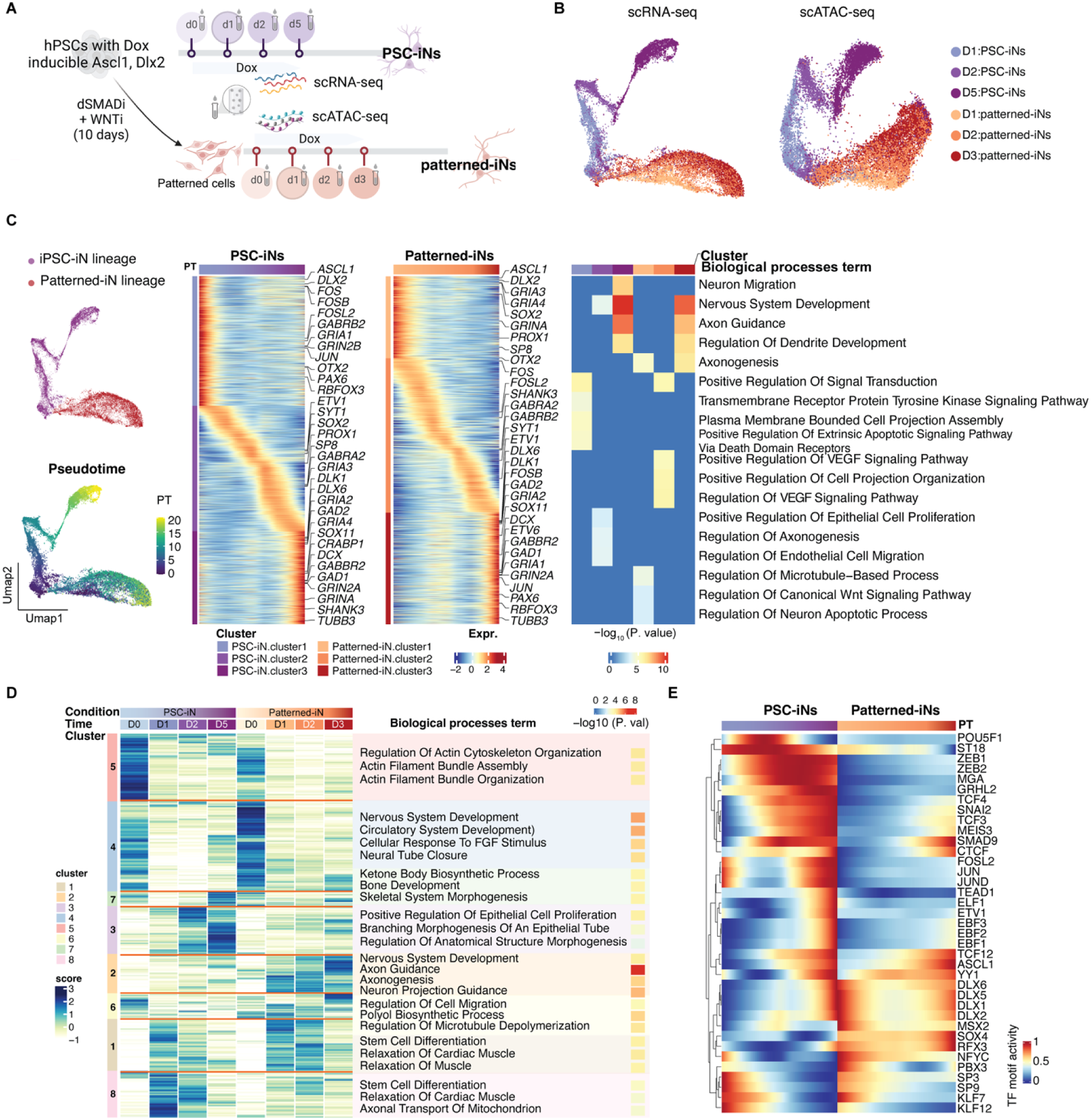
Molecular Signatures Underlying iN Specification. **(A)** Overview of the experimental workflow. **(B)** UMAP representations based on gene expression (left) and chromatin accessibility (right). Cells are color-coded according to differentiation time points and methods used. **(C)** UMAP of single-cell RNA sequencing (scRNA-seq) data highlighting the differentiation trajectory of PSC-iNs and patterned iNs, along with Slingshot-derived pseudotime (left). The middle panel shows a heatmap of gene expression across pseudotime bins within the PSC-iNs and patterned iNs trajectory. The right panel presents the Gene Ontology (GO) analysis of genes induced at various stages during early differentiation. **(D)** Heatmap showing the chromatin accessibility across pseudobulk samples, aggregated from individual differentiation stages along the PSC-iNs and patterned iNs trajectory. The right panel shows the GO analysis of genes associated with accessible chromatin regions across the eight clusters. **(E)** Heatmaps showing motif activity of TFs across pseudotime bins. See also Figures S8 and S9.

To investigate the mechanisms governing the specification of GABAergic neurons derived from PSCs and patterned progenitors, we first reconstructed the trajectory of neuron development by assigning pseudotime values to individual cells and tracing their progression along their respective differentiation pathways (Figure 5C). As expected, cells treated using small molecules exhibited expression of neural progenitor cell (NPC) marker genes (e.g., *SOX2*, *NES*, *SOX11*, *PAX6*) (Figure S8). Following TF induction, genes associated with interneuron development, such as *OTX2*, *PROX1*, *TCF12*, and *RELN*, were robustly induced in the patterned progenitors and, to a lesser extent, in iPSCs (Figures 5C and S8). This expression pattern quickly transitioned towards a GABAergic neuronal identity, marked by the induction of *GAD1*, *GAD2*, *DLX6*, and related markers. In parallel, neuronal differentiation genes (e.g., *DCX, TUBB3, RBFOX3*, etc.) and genes encoding neurotransmitter receptors (*GRIA1-4*, *GRINA*, *GRIN2A*, *GABRA2/B2*, etc.) were induced, consistent with the rapid gene expression changes characteristic of TF-mediated cell fate conversion. To complement these analyses and assess lineage specificity, we examined canonical markers of excitatory neuronal and glial lineages in both PSC-iNs and patterned-iNs. Markers of cortical excitatory neurons (*TBR1*, *EMX1*, *SLC17A6/7*) were undetectable across iGABAs. Similarly, astrocytic (*GFAP*, *AQP4*, *S100B*, *ALDH1L1*) and oligodendrocytic (*OLIG1*, *OLIG2*, *MBP*, *MOG*) markers were absent or expressed at negligible levels (Figure S9). *MBP* showed modest induction in PSC-iNs but not in patterned-iNs, suggesting a slightly higher propensity for glial gene activation in PSC-derived cells. Together, these data confirm efficient and selective induction of inhibitory neuronal identity by ASCL1 and DLX2 in both contexts, while revealing subtle differences in lineage restriction between differentiation strategies.

Genes active at later pseudotime stages were enriched for gene ontology (GO) terms related to neural morphogenesis, migration, and development (Figure 5C). In contrast, distinctive GO enrichments were associated with genes active at early and intermediate stages of differentiation trajectories over pseudotime, suggesting that ASCL1 and DLX2 induce context-dependent molecular programs in PSCs and patterned progenitors. Specifically, GO terms uniquely enriched among genes active early in the PSC-iN trajectory included non-neuronal processes, whereas genes induced in patterned iNs were associated with signaling pathways involved in neurogenesis, including WNT and VEGF signaling pathways ^68–70^.

Further analysis of the chromatin landscapes before and after TF induction revealed significant state-dependent remodeling, with distinct clusters of open chromatin peaks emerging across different cell states and stages of differentiation (Figure 5D). GO analysis of genes linked to these open chromatin regions showed that regions accessible in PSCs and patterned NPC-like cells but rendered largely inaccessible after ASCL1 and DLX2 induction were primarily associated with non-neuronal processes (clusters 4, 5). Conversely, regions that were initially inaccessible in PSCs or patterned cells but became accessible after TF induction were strongly enriched for neurogenesis (cluster 2). We also noted that transiently accessible regions were linked to stem cell differentiation (clusters 1, 8), whereas regions that became robustly accessible exclusively along the PSC-iN trajectory were occasionally associated with “off-target” regulator programs (clusters 3, 7). Together, these chromatin dynamics mirror the transcriptional changes observed following TF induction and indicate a more stringent and lineage-directed neuronal differentiation program in patterned NPC-like cells, compared with a more heterogeneous regulatory program in PSCs at early stages. These findings highlight the nuanced interplay between epigenetic and transcriptional regulation in shaping cell fate decisions.

We next examined TF-driven regulatory mechanisms across pseudotime by correlating TF expression with motif activity scores derived from chromatin accessibility. This analysis identified 37 TF motifs differentially associated with the progression along the PSC-iN or patterned-iN differentiation trajectories (Figure 5E). Both differentiation pathways showed strong activity of ASCL1 and DLX motifs, reinforcing their central roles in cell-fate determination. Additionally, motifs corresponding to TFs with established roles in neural development and function, including TCF12^71^, CTCF^72,73^, SP3^74^, SP9^75^, and KLF7^76,77^, were active in both differentiation trajectories. In contrast, numerous other TF motifs, particularly those integral to pluripotency maintenance and broader lineage plasticity, such as POU4F1 (OCT4), MGA^78^, SNAI2^79^, TCF3^80^, and MEIS3^81^ were more specifically involved in the PSC-iN differentiation pathway. Collectively, these findings underscore the intricate coordination of TF motifs early in neuronal differentiation, supporting our observation that ASCL1 and DLX2 engage a more tightly constrained TF network in patterned progenitors, fostering a more defined path toward GABAergic neuronal differentiation. In contrast, PSCs retain greater plasticity, enabling activation of a broader spectrum of TF-driven gene expression networks. This distinction suggests that although both PSCs and patterned progenitors are competent for neuronal differentiation, the pathways and regulatory networks involved are significantly divergent, reflecting the intrinsic differences in their biological potential and responsiveness to transcriptional cues.

### Disease Modeling Potential of Patterned Induced GABAergic Neurons

To evaluate whether GABAergic patterned-iNs can be leveraged to uncover disease-relevant phenotypes, we focused on ADNP syndrome, a rare neurodevelopmental disorder caused by mutations in the *activity-dependent neuroprotective protein (ADNP)* gene and frequently associated with ASD. ^82–84^ GABAergic interneurons are central to regulating excitation–inhibition balance and network synchrony, processes commonly disrupted in ASD and related NDDs, yet the effects of ADNP mutations on human inhibitory neurons have not been well characterized.

Among individuals with ADNP syndrome, a recurrent heterozygous nonsense mutation at p.Tyr719* has been frequently identified.^82–84^ To model this variant, we used CRISPR-Cas9 to introduce the ADNP p.Tyr719* allele into two independent human iPSC lines (PGP1 and WTC11). In parallel, a hemagglutinin (HA) tag was inserted at the 3’ end of the endogenous *ADNP* locus or immediately following the p.Tyr719* truncation to enable detection of full-length and truncated ADNP proteins. As shown in Figure S10, an ADNP antibody recognizing a region between residues 1,050 and the C-terminus detected full-length ADNP in both unedited and HA-tagged control cells but not the truncated p.Tyr719* protein, whereas the truncated protein was readily detected using an HA antibody. Consistent with heterozygosity, levels of full-length ADNP were reduced in heterozygous p.Tyr719* mutant lines relative to unedited controls.

Patterned-iNs were generated from heterozygous and homozygous *ADNP* p.Tyr719* iPSCs and their corresponding isogenic controls (six lines in total) and analyzed five weeks after differentiation in glial co-culture by scRNA-seq. Following quality control and filtering, the WTC11 homozygous cells were excluded from subsequent analyses due to insufficient cell recovery. Patterned iNs derived from different genetic backgrounds exhibited high reproducibility (Figure S11A). Transcriptomic profiling revealed a high degree of correlation between heterozygous and homozygous mutant cells (Figures 6A and 6B), prompting us to combine mutant cells from both PGP1 and WTC11 lines to increase statistical power. Pan-neuronal markers, including MAP2 and RBFOX3 (NeuN), were robustly expressed in both wild-type and mutant neurons (Figure S11B), confirming successful neuronal differentiation. Assessment of neurotransmitter identity showed that the majority of neurons were GABAergic, with 98.62% of wild-type and 97.28% of mutant MAP2-positive neurons classified as GABAergic.

**Figure 6.**
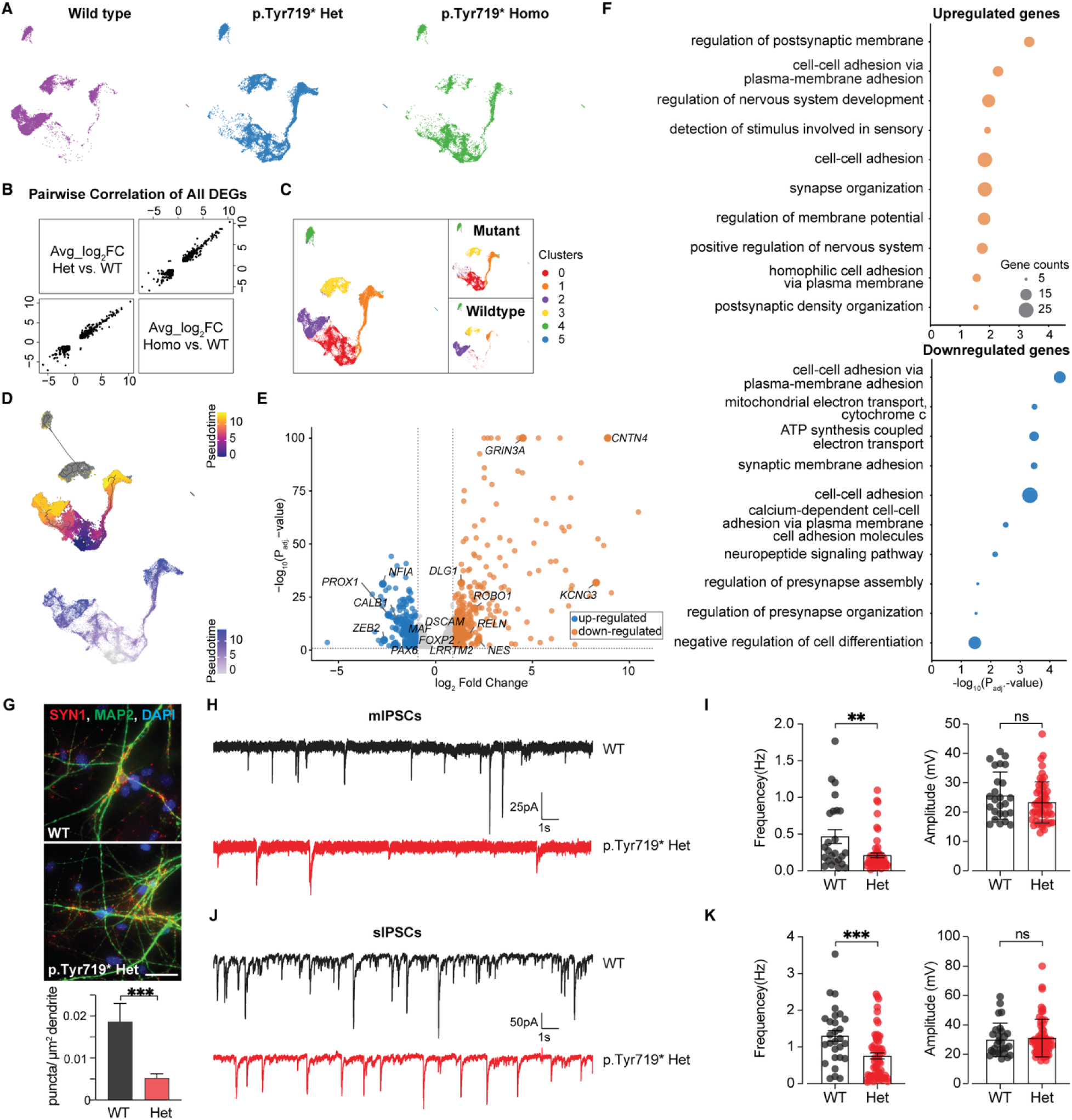
ADNP p.Tyr719 Mutation Disrupts Neurogenic and Synaptic Gene Expression and Function. (**A**) UMAP embedding of cells grouped by ADNP genotype (wild-type, heterozygous, and homozygous) across different iPSC lines, showing clustering by transcriptional similarity. (**B**) Pairwise correlation heatmap of all differentially expressed genes (DEGs) between wild-type and ADNPp.Tyr719* samples, indicating similar transcriptional changes caused by both heterozygous and homozygous ADNPp.Tyr719* mutations. (**C**) Combined UMAP of all single-cell transcriptomes reveals genotype-enriched clusters. Right: Same UMAP split by genotype, highlighting differences in cell distribution between WT and ADNP-mutant cells. (**D**) Pseudotime UMAP analysis of all cell types (top), and pseudotime value of clusters 0, 1, and 2 identified in (C) (bottom). (**E**) Volcano plot displaying DEGs between ADNPp.Tyr719* and WT neurons (adjusted p < 0.05; |log₂FC| > 1). Selected upregulated and downregulated genes are annotated. (**F**) Top Gene Ontology (GO) Biological Process terms enriched among DEGs shown in (E), highlighting disruption of neurodevelopmental and synaptic programs. (**G**) Immunofluorescence images of SYN1 (synapsin 1), MAP2, and DAPI staining in WT and ADNP Het neurons. Right: Quantification of SYN1 puncta density per μm² of dendrite, revealing reduced synaptic density in ADNP Het neurons. Scale bar: 50 μm. (**H**) Representative traces of miniature inhibitory postsynaptic currents (mIPSCs) recorded from WT (blue) and ADNPp.Tyr719* heterozygous (Het, red) neurons. (**I**) Quantification of mIPSC frequency and amplitude from recordings in (**H**), across WT (blue, n = 25) and ADNP Het neurons (n = 61 from three batches). (**J**) Representative traces of spontaneous inhibitory postsynaptic currents (sIPSCs) recorded from WT and ADNP Het neurons. (**K**) Summary of sIPSC frequency and amplitude across pooled WT (n = 28 from three batches) and ADNP Het neurons (n = 61 from three batches), demonstrating significantly reduced inhibitory synaptic input in ADNP mutant cells. Data are shown as mean ± SEM. Statistical comparisons were made using one-way ANOVA (for multi-group) or unpaired t-tests (for two-group). *P < 0.05; **P < 0.01; ***P < 0.001. See also Figures S10, S11, S12 and Tables S5, S6, S7, S8, S9 and S10.

To assess cellular heterogeneity, we applied the Leiden clustering algorithm to identify transcriptomically distinct populations. This analysis revealed clear segregation between mutant and wild-type neurons, supported by statistically significant differences in gene expression (Table S5). Specifically, clusters 0 and 1 were predominantly composed of p.Tyr719* mutant cells, whereas cluster 2 was enriched for wild-type neurons (Figure 6C). We next examined maturation states by assigning pseudotime values using Monocle3, which identified cluster 0 mutant neurons as the least mature population (Figure 6D). This interpretation was supported by enrichment of *NESTIN* (*NES*), a marker commonly associated with neural progenitors, within this cluster (Table S5). A continuous differentiation trajectory within the mutant population was also observed and independently validated using Slingshot (Figure S11C). Along this trajectory, mutant neurons progressively upregulated genes encoding GABA and glutamate receptor subunits (*GABRB2*, *GRIA2*, *GRID2*), synaptic signaling components (*SYN3*, *DLG2*, *NRXN3*), and neuronal differentiation markers (*ROBO1*, *NRP2*), all of which are critical for neuronal maturation and synaptic transmission (Figure S11D; Table S6).

To further define molecular alterations associated with the p.Tyr719* mutation, we examined DEGs between mature mutant and wild-type cells using a pseudobulk analysis framework. Mutant neurons exhibited both downregulated and upregulated transcriptional programs (Figure 6E and Table S7). Notably, genes involved in the generation of CGE-derived cortical interneurons, including *NR2F2* and *PROX1*, as well as genes associated with LGE-derived interneurons in the olfactory bulb and basal nuclei, such as *FOXP2,* and *CALB1*,^13^ were significantly downregulated in the mutant cells. PAX6^13,49^, which contributes to both CGE and LGE interneuron development, was also reduced. Conversely, *RELN*, primarily produced by specific subsets of GABAergic interneurons in the neocortex and hippocampus, was upregulated.^85,86^ GO analysis of DEGs highlighted enrichment for pathways related to synaptic organization and neuronal connectivity, including regulation of postsynaptic membrane potential, synapse organization, trans-synaptic signaling, and cell–cell adhesion, as well as processes involved in neuronal development and maturation, such as regulation of neuron projection development and nervous system development (Figure 6F and Table S8).

To capture coordinated transcriptional changes associated with the p.Tyr719* mutation, we performed Weighted Gene Co-expression Network Analysis (WGCNA) focusing on clusters 0, 1, and 2. To mitigate the sparsity inherent in single-cell data while preserving the heterogeneity often masked by traditional cell clustering, we first constructed metacells^87^, representing aggregates of cells with similar gene expression profiles, in which within-metacells variation is primarily technical rather than biological (Figure S12A). Subsequently, we constructed gene co-expression modules separately for the three clusters, identifying 25 gene co-expression modules (Figure S12B and Table S9). Modules associated with mutant neurons were enriched for biological processes including translational initiation, neuronal development, membrane potential regulation, and chemical synaptic transmission (Figure S12C).

Finally, to determine whether these transcriptional alterations translated into functional deficits, we performed whole-cell patch-clamp recordings on patterned GABAergic interneurons derived from three independent ADNP p.Tyr719* mutant clones and matched controls. Mutant neurons exhibited significantly reduced spontaneous and miniature inhibitory postsynaptic current (sIPSC and mIPSC) frequencies, indicating impaired inhibitory synaptic transmission (Figures 6G-K and Table S10). These deficits were consistent across three independent differentiation batches, demonstrating robustness across genetic backgrounds and experimental replicates. Collectively, these results show that the ADNP p.Tyr719* mutation disrupts interneuron maturation, subtype-associated gene expression, and inhibitory synaptic function in human patterned GABAergic neurons, supporting the utility of patterned-iNs as a platform for identifying mutation-specific mechanisms in neurodevelopmental disease.

## DISCUSSION

Differentiating human stem cells into regional- and neurotransmitter-specific subtypes of human interneuron populations offers a promising platform for studying pathophysiology and potential therapeutic approaches for various diseases. This study presents a rapid and reproducible method for generating GABAergic neurons by integrating our TF-mediated PSC differentiation^27^ with developmental patterning achieved by inhibiting SMAD and WNT signaling^36,42^. This method, akin to the direct conversion of PSCs into GABAergic neurons driven by ASCL1 and DLX2, yields a homogeneous population of cells with inhibitory identity. We provide an extensive, mineable single-cell transcriptomic resource encompassing GABAergic neurons generated using TF-based induction, small-molecule-only patterning, and combinatorial patterning plus TF approaches. Systematic comparison across these differentiation strategies reveals that each method yields GABAergic neurons with overlapping yet distinct molecular identities, developmental trajectories, and subtype compositions. Importantly, this diversity is not a limitation but a strength for disease modeling, as different neurodevelopmental and neuropsychiatric disorders are associated with dysfunction in certain interneuron subtypes and developmental programs. By enabling direct cross-method benchmarking, this dataset facilitates informed selection of PSC-derived GABAergic neuron models best suited for interrogating particular genes, pathways, or disease contexts. Additionally, the study demonstrates that this combinatorial approach produces disease-relevant GABAergic neuron types and applies it to investigate the pathobiology of ADNP syndrome.

SMAD and WNT inhibitors can induce forebrain lineages from human PSCs, including excitatory and inhibitory neurons.^36,42^ By activating SHH signaling, progenitor cells can be ventralized to differentiate toward human cortical interneuron lineages.^42,43,45,56^ However, the maturation of these progenitor cells requires a protracted time, as evidenced by scRNA-seq data. When ASCL1 and DLX2 are induced in PSCs exposed to SMAD and WNT inhibitors, homogeneous GABAergic neurons can be derived without the need for SHH or its agonist. The neurons derived under this condition represent multiple brain regions and are biased toward the cortical lineage. In contrast, PSC-iGABAs without patterning molecules exhibit a more prominent hypothalamic identity. Furthermore, the PSC-iGABAs are enriched for TF regulons associated with hypothalamic and striatal neurogenesis and function. On the other hand, patterned-iGABAs are enriched for TF regulons linked to neural fate determination and migration. This difference highlights the distinct developmental trajectories and functional specializations of the two types of iGABAs. Importantly, when transplanted into the neonatal mouse brain, patterned iGABAs survive long term and differentiate into mature neurons but remain largely localized near the injection site, a limited migration pattern consistent with prior reports of PSC-iGABAs^27^ and likely reflecting direct, postmitotic conversion driven by ASCL1 and DLX2 rather than expansion of migratory neuroblast populations. Notably, recent work utilizing barcoding tools for lineage tracing suggested that individual human cortical progenitors can produce both excitatory and inhibitory neurons.^88^ Cortically-derived inhibitory neurons share transcriptional similarities with CGE-derived interneurons but lack definitive marker genes. Given the expression of *SP8* and *NR2F2* in the patterned iNs, it is possible that some of these neurons represent cortically-derived inhibitory neurons.

The differential gene expression programs observed in PSC-iGABAs and patterned-iGABAs generated through ASCL1 and DLX2 overexpression highlight the complexity of transcriptional regulation in neurogenesis. Multiple factors may contribute to these differences. One of the primary reasons is the differences between the epigenetic landscapes of PSCs and NPC-like cells obtained by SMAD and WNT inhibition. PSCs possess a more open and plastic chromatin structure, allowing a broader range of TF binding sites to be accessible. In contrast, NPCs have a more restricted chromatin landscape, with specific regions pre-configured for neural differentiation. ASCL1, as a pioneering factor, can bind to closed chromatin regions and initiate chromatin remodeling. However, the function of pioneering factors is heavily influenced by cellular context. In PSCs, ASCL1 can access and activate a wide array of genes, leading to diverse gene expression profiles that may even include non-neuronal lineages. Conversely, the chromatin landscape in the NPCs is already more accessible in regions relevant to neural development, so ASCL1 and DLX2 primarily enhance the existing neural gene expression program. PSCs and NPCs also differ in their baseline transcriptional states. PSCs express genes associated with pluripotency and self-renewal, whereas NPCs are committed to a neural lineage and express genes characteristic of neural progenitor cells. When ASCL1 and DLX2 are introduced into these different contexts, the downstream targets they activate are influenced by the pre-existing transcriptional milieu. ASCL1 and DLX2 might interact with pluripotency factors and other non-neural transcriptional networks, leading to a more heterogeneous response in PSCs. Meanwhile, these TFs reinforce and enhance the neural differentiation program in NPCs, leading to a more focused and streamlined gene expression profile. This aligns with the observation that GABAergic patterned-iN subtypes are less molecularly distinguishable. Additionally, the availability of cofactors and interaction partners in PSCs and NPC-like cells can further modulate the activity of ASCL1 and DLX2, contributing to the observed differences in gene expression programs.

Finally, we leveraged patterned human GABAergic interneurons to investigate the phenotypic consequences of the ADNP syndrome-associated p.Tyr719* variant. ADNP is ubiquitously expressed and functions as a chromatin-associated regulator of gene expression through interactions with multiple partners, including CHD4, a core component of the nucleosome remodeling and deacetylase (NuRD) complex.^89,90^ Together with CHD4 and HP1γ, ADNP forms the ChAHP complex, which represses transcription at specific genomic loci.^89^ In *Adnp* knockout embryonic stem (ES) cells and mouse embryos, defects in neuronal fate specification have been observed,^89^ and impaired excitatory and inhibitory synapses have been reported in juvenile *Adnp* haploinsufficient mice.^91^ However, most existing *in vivo* studies rely on large exonic deletions that do not reflect the molecular nature of pathogenic human ADNP mutations. Large-scale genetic studies of ASD and developmental delay disorders have revealed a striking enrichment of *de novo* protein-truncating variants in *ADNP*, with a strong bias toward mutations located in the terminal exon, including the highly recurrent p.Tyr719* variant. These mutations are predicted to escape nonsense-mediated decay, and mutant *ADNP* transcripts have been detected in patient-derived samples, including peripheral blood, suggesting that truncated proteins are expressed. ^84,92^ Consistent with this prediction, we used endogenous HA tagging to directly demonstrate stable expression of the truncated ADNP protein from the native locus in human cells, providing direct evidence that pathogenic ADNP truncations are not simple loss-of-function alleles. This observation supports the possibility that disease-causing *ADNP* mutations may exert dominant-negative or gain-of-function effects, a mechanistic distinction that is not captured by deletion-based models.

Consistent with this interpretation, our data suggest that the p.Tyr719* variant not only alters the specification of neuronal subtypes within the GABAergic lineage but may also disrupt synaptic transmission in differentiated cells. Notably, we observed a high degree of transcriptional similarity between heterozygous and homozygous p.Tyr719* mutant interneurons, in contrast to prior reports showing minimal transcriptional differences between wild-type and heterozygous conditional Adnp knockout animals.^89^ One plausible explanation is that, unlike the removal of exon 5, which encodes all but 67 of Adnp’s 1108 total amino acids, the truncated ADNP protein produced by p.Tyr719* competes with full-length ADNP for chromatin binding sites or interacting partners, thereby perturbing gene regulatory networks in a manner distinct from simple dosage reduction.^89^ Intriguingly, we observed upregulation of *CHD4* expression in mutant cells, a phenomenon not reported in *Adnp* knockout models (Figure S10B). Given that p.Tyr719* interacts with CHD4, this upregulation may reflect compensatory responses to the sequestration of CHD4 by p.Tyr719*. Further studies will be required to define the precise molecular mechanisms underlying these effects.

Overall, our study expands the toolbox for disease modeling by demonstrating the complementary strengths of PSC-iNs and patterned iNs in representing different brain regions and disease contexts. This dual approach offers a robust platform for investigating the molecular mechanisms underlying a variety of neurodevelopmental and neuropsychiatric disorders.

## Acknowledgments

This work was supported by NIH awards 5R21MH119580 (N.Y.), 5R01NS116057 (N.Y.), R21MH131600 (J.D.B., N.Y.), 5R01AA023797 (Z.P.P), and 5R01MH125528 (Z.P.P). This study was supported in part by a grant from the Simons Foundation (SFI-AN-AR-Pilot-00009949) (J.D.B.), Oxford-Harrington Rare Disease Scholar Award (J.D.B.), ADNP Foundation, and Seaver Foundation. X. W. is supported as a Seaver Foundation Fellow and by NARSAD Young Investigator grants (31904). R.H. is a Seaver Postdoctoral Fellow. L.L.B. was supported by the Training Program in Stem Cell Biology fellowship from the New York State Department of Health (NYSTEM-C32561GG). M.Y. is supported by a diversity supplement (3R01MH129372-03S1). M.L. is supported by the NIH F31 Ruth L. Kirschstein National Research Service Award (F31NS137751).

## Author contributions

Conceptualization, R.H., L.L.B., J.D.B., S.M., and N.Y.; Methodology, R.H., L.L.B., and Y.T.; Investigation, R.H., Y.T., B.Z., J.L., X.L., R.D., M.L., G.I.G., and N.Y.; Formal Analysis and Data Visualization, L.L.B., M. Y., P.C., Y.S., J.L., X.L., X.W., A.T., and S.M.; Writing, R.H., L.L.B., J.L., X.L., X.W., S.M., and N.Y.; Supervision, X. W., A.T., J.D.B., P.J., Z.P.P., S.M., and N.Y.

## STAR Methods

Detailed methods are provided in the online version of this paper and include the following:

**KEY RESOURCES TABLE**
**RESOURCE AVAILABILITY**
`o` Lead contact
`o` Materials availability
`o` Data and code availability
EXPERIMENTAL MODEL AND SUBJECT DETAILS

`o` Animal Model
`o` Human PSC lines and culture
`o` Lentivirus production
`o` Small molecule-mediated GABAergic neuron generation
`o` TF-mediated induced GABAergic neuron generation
`o` Patterned induced GABAergic neuron generation
`o` Lentivirus production
`o` Lentivirus transduction
**METHOD DETAILS**

`o` Primary mouse astrocytes
`o` Transplantation and brain slice preparation
`o` Electrophysiological recordings from brain slices
`o` Whole-cell recordings from cultured neurons
`o` Immunofluorescence and imaging
`o` Immunoblotting
`o` 10X single-cell RNA-seq library preparation and sequencing
`o` SHARE-seq library preparation and sequencing
**QUANTIFICATION AND STATISTICAL ANALYSIS**

`o` 10X Single-cell RNA-sequencing data processing and analysis
`o` SHARE-seq data processing and analysis
`o` Inferring regulatory networks
`o` Gene Ontology, pathway enrichment, and disease-vulnerable cell type
`o` WGCNA analysis
`o` Statistical analyses

**STAR Methods**

### KEY RESOURCES TABLE

| REAGENT or RESOURCE | SOURCE | IDENTIFIER |
| --- | --- | --- |
| Antibodies |  |  |
| Anti-ADNP | Bethyl Laboratories, Inc | Cat # A300-104A; RRID:AB_242522 |
| Anti-HA | Roche | Cat #11867423001;RRID:AB_390918 |
| Anti-CTCF | Cell Signaling Technology | Cat # 3418S; RRID:AB_2086791 |
| Anti-HP1γ | Cell Signaling Technology | Cat # 2619S; RRID:AB_2070984 |
| Anti-CHD4 | Bethyl Laboratories, Inc | Cat # A301-081A; RRID:AB_873001 |
| Anti-MAP2 | Abcam | Cat # ab5392; RRID:AB_2138153 |
| Anti-Synapsin | Synaptic Systems | Cat # 106 002; RRID:AB_887804 |
| Anti-FOXG1 | Takara Bio | Cat # M227; RRID:AB_2827749 |
| Anti-Sox2 | R&D Systems, Inc. | Cat # AF2018; RRID:AB_355110 |
| Anti-Pax6 | BioLegend | Cat # 901301; RRID:AB_2565003 |
| Anti-H3K27ac | Abcam | Cat # AB4729; RRID:AB_2118291 |
| Anti-IgG | Abcam | Cat # AB46540; RRID:AB_2614925 |
| Anti-IgG | Novus Biologicals, LLC | Cat # NBP1-2763 |
| Anti-FOS | Abcam | Cat # AB208942; RRID:AB_2747772 |
| Anti-GAPDH | Cell Signaling Technology | Cat # 2118; RRID:AB_561053 |
| Anti-β-actin | Sigma-Aldrich | Cat # A5441p RRID:AB_476744 |
| Anti-Dlx5 | (Wang et al., 2010) | Gift from John LR Rubenstein. |
| Anti-Dlx2 | Santa Cruz Biotechnology, Inc | Cat # sc-393879 |
| Anti-GAD67 | Millipore | Cat# MAB5406, RRID:AB_2278725 |
| Anti-CALB2 | Agilent | Cat# M7245, RRID:AB_2068519 |
| Anti-Calbindin | Swant | Cat# 300, RRID:AB_10000347 |
| Anti-Pax6 | Covance | Cat# PRB-278P, RRID:AB_291612 |
| Anti-Somatostatin | Novus | Cat# NBP2-37447, RRID:AB_2890053 |
| Anti-Human Nuclear Antigen | Millipore | Cat# MAB1281, RRID:AB_94090 |
| Anti-SP8 | Santa Cruz Biotechnology | Cat# sc-104661, RRID:AB_2194626 |
| Anti-Synaptophysin | Thermo Fisher Scientific | Catalog # 14-6525-82; RRID:AB_10670280 |
| Anti-NeuN | Millipore | Cat# MAB377, RRID:AB_2298772 |
| Peroxidase-AffiniPure Donkey Anti-Rabbit IgG | Jackson ImmunoResearch Labs | Cat# 711-035-152, RRID:AB_10015282 |
| IRDye 680RD Donkey anti-Mouse IgG | LI-COR Biosciences | Cat# 926-68072, RRID:AB_10953628 |
| IRDye 800CW Goat anti-Rat IgG | LI-COR Biosciences | Cat# 926-32219, RRID:AB_1850025 |
| Donkey anti-Mouse IgG 488 | Thermo Fisher Scientific | Cat# A-21202, RRID:AB_141607 |
| Donkey anti-Mouse IgG 555 | Thermo Fisher Scientific | Cat# A-31570, RRID:AB_2536180 |
| Donkey anti-Rabbit IgG 555 | Thermo Fisher Scientific | Cat# A-31572, RRID:AB_162543 |
| Donkey anti-Rabbit IgG 488 | Thermo Fisher Scientific | Cat# A-21206, RRID:AB_2535792 |
| Donkey anti-Chicken IgG 647 | Thermo Fisher Scientific | Cat# A78952, RRID:AB_2921074 |
| Chemicals, peptides, and recombinant proteins |  |  |
| Stemflex supplement | Gibco \| Thermo Fisher Scientific | Cat # A33492 |
| Stemflex basal media | Gibco \| Thermo Fisher Scientific | Cat # A33493-01 |
| MEM media | Gibco \| Thermo Fisher Scientific | Cat # 11090-081 |
| Neurobasal media | Gibco \| Thermo Fisher Scientific | Cat # 21103-049 |
| Neurobasal A media | Gibco \| Thermo Fisher Scientific | Cat # 10888-022 |
| DMEM/F12 media | Gibco \| Thermo Fisher Scientific | Cat # 11320-033 |
| DMEM media | Gibco \| Thermo Fisher Scientific | Cat # 11965-092 |
| Knockout DMEM | Gibco \| Thermo Fisher Scientific | Cat # 10829018 |
| B27 supplement | Gibco \| Thermo Fisher Scientific | Cat # 17504-044 |
| N2 supplement | Gibco \| Thermo Fisher Scientific | Cat # 17502-048 |
| KnockOut™ Serum Replacement | Gibco \| Thermo Fisher Scientific | Cat # 10828028 |
| MEM NEAA (100x) | Gibco \| Thermo Fisher Scientific | Cat # 11140-050 |
| Glutamax (100x) | Gibco \| Thermo Fisher Scientific | Cat # 35050-061 |
| Sodium Pyruvate (100x) | Gibco \| Thermo Fisher Scientific | Cat # 11360-070 |
| β-Mercaptoethanol | Sigma-Aldrich | Cat # M3148 |
| Cosmic Calf™ Serum | Cytiva | Cat # SH3008704 |
| Fetal Bovine Serum | Sigma-Aldrich | Cat # F6178 |
| D-PBS tablets | Millipore | Cat # 524650-1EA |
| Geltrex | Gibco \| Thermo Fisher Scientific | Cat # A14133-02 |
| Poly-L-ornithine hydrobromide | Sigma-Aldrich | Cat # P3655 |
| Accutase | Innovative Cell Technologies | Cat # AT104500 |
| Trypsin 0.25% | Gibco \| Thermo Fisher Scientific | Cat # 25200-114 |
| Puromycin dihydrochloride | Sigma-Aldrich | Cat # P8833 |
| Hygromycin B (50 mg/mL) | Gibco \| Thermo Fisher Scientific | Cat # 10687010 |
| Cytosine β-D-arabinofuranoside | Sigma-Aldrich | Cat # C1768 |
| Doxycycline | Sigma-Aldrich | Cat # P3655 |
| Bovine Serum Albumin | Equitech-Bio Inc | Cat # BAH65-0500 |
| Thiazovivin | Santa Cruz Biotechnology, Inc | Cat # SC-361380 |
| Y-27632 | Tocris Bioscience | Cat # 1254 |
| Chroman1 | MedChemExpress LLC | Cat # HY-15392 |
| LDN193189 | Stemgent | Cat # 04-0074 |
| SB431542 | Tocris Bioscience | Cat # 1614 |
| XAV939 | Stemgent | Cat # 04-0046 |
| DAPT | Tocris Bioscience | Cat # 2634 |
| SHH | R&D Systems | Cat # 1845-SH |
| Purmorphamine | R&D Systems | Cat # SML0868 |
| Dimethyl sulfoxide | Sigma-Aldrich | Cat # D2438 |
| Paraformaldehyde solution | Affymetrix | Cat # 19943 |
| Halt Protease and<br>Phosphatase Inhibitor Cocktail | Thermo Fisher Scientific | Cat # 78440 |
| Critical commercial assays |  |  |
| Pierce RIPA Buffer | Thermo Fisher Scientific | Cat # 89900 |
| Bolt™ 4 to 12%, Bis-Tris,<br>Protein Gel | Thermo Fisher Scientific | Cat # MW04125BOX |
| iBlot™ 2 Transfer Stacks,<br>PVDF | Thermo Fisher Scientific | Cat # IB24001 |
| Western Lightning Plus,<br>Chemiluminescent Substrate | Revvity Health Sciences Inc | Cat # NEL104001EA |
| Pierce™ BCA Protein Assay<br>Kits | Thermo Fisher Scientific | Cat # 23225 |
| Intercept® (PBS) Blocking<br>Buffer | LI-COR bio | Cat # P/N: 927-70001 |
| Bio Rad ChemiDoc MP<br>Imaging System | Bio Rad | RRID:SCR_019037 |
| Chromium Next GEM Single<br>Cell 3' GEM Kit v3.1 | 10X Genomics | Cat # PN-1000123 |
| Chromium Next GEM Single<br>Cell 3' Gel Bead Kit v3.1 | 10X Genomics | Cat # PN-1000122 |
| Chromium Next GEM Chip G<br>Single Cell Kit | 10X Genomics | Cat # PN-1000120 |
| Chromium iX | 10X Genomics | Cat # PN-1000328,<br>RRID:SCR_024536 |
| ZEISS LSM 980 with Airyscan<br>2 Microscope | ZEISS | RRID:SCR_025048 |
| EVOS M5000 Imaging System | ThermoFisher Scientific | Cat # AMF5000,<br>RRID:SCR_023650 |
| Deposited Data |  |  |
| Raw single cell-sequencing<br>data and normalized counts | This paper | GSE277602 and GSE277604 |
| Experimental models: cell lines |  |  |
| PGP1 | Coriell | Cat# GM23338, RRID:CVCL_F182 |
| WTC11 | Coriell | Cat# GM25256, RRID:CVCL_Y803 |
| HES-3 NKX2.1GFP/w | Goulburn A.L., et al., 2011<sup>93</sup> | RRID:CVCL_A5HB |
| HEK293T | ATCC | Cat # 11268, RRID:CVCL_1926 |
| Experimental models: organisms/strains |  |  |
| CD1 mouse strain | Charles River | Cat# CRL:022,<br>RRID:IMSR_CRL:022 |
| Recombinant DNA |  |  |
| FUW-M2rtTA | Hockemeyer et al., 2008<sup>94</sup> | RRID:Addgene_20342 |
| TetO-Ascl1-puro | Yang et al., 2017<sup>27</sup> | RRID:Addgene_97329 |
| TeoO-Dlx2-Hygro | Yang et al., 2017<sup>27</sup> | RRID:Addgene_97330 |
| pMD2.G | Addgene | RRID:Addgene_12259 |
| pRSV-rev | Addgene | RRID:Addgene_12253 |
| pMDLg/pRRE | Addgene | RRID:Addgene_12251 |
| PB-TO-ASCL1-DLX2 | Addgene | RRID:Addgene_182307 |
| Software and algorithms |  |  |
| Cellranger v7.2.0 | 10x Genomics | RRID:SCR_023221 |
| ClusterProfiler v4.8.3 | Yu et al., 2012<sup>95</sup> | RRID:SCR_016884 |
| DoubletFinder v2.0.4 | McGinnis et al., 2019<sup>96</sup> | RRID:SCR_018771 |
| Gephi v0.10.1 | http://gephi.org/ | RRID:SCR_004293 |
| ggplot2 v3.4.3 | Wickham, 2016<sup>97</sup> | RRID:SCR_014601 |
| Metacell | Baran et al., 2019<sup>87</sup> | RRID:SCR_017013 |
| Monocle3 | Cao et al., 2019<sup>98</sup> | RRID:SCR_018685 |
| ZEISS ZEN Microscopy<br>Software | ZEISS | RRID:SCR_013672 |
| R | Love et al., 2014<sup>99</sup> | https://www.r-project.org/ |
| Revigo v1.8.1 | Supek et al., 2011<sup>100</sup> | RRID:SCR_005825 |
| RStudio v4.3.1 | Rstudio team, 2020<sup>101</sup> | https://github.com/rstudio/ |
| Seurat v4.0.1. | Butler et al., 2018<sup>102</sup> | RRID:SCR_016341 |
| SingleCellExperiment v1.22.0 | Amezquita et al., 2020<sup>103</sup> | https://github.com/drisso/SingleCellExperiment |
| Slingshot v2.8.0 | Street et al., 2018<sup>104</sup> | RRID:SCR_017012 |

### RESOURCE AVAILABILITY

#### Lead contact

Further information and requests for resources and reagents should be directed to and will be fulfilled by the Lead Contact, Nan Yang.

#### Materials availability

The plasmids and cell lines generated in this study will be made available on request upon the completion of a Material Transfer Agreement (MTA).

#### Data and code availability

Single-cell RNA-seq and SHARE-seq data have been deposited at GEO and are publicly available as of the date of publication. Accession numbers are listed in the key resources table. The DOI is listed in the key resources table. This paper does not report original codes. Any additional information required to reanalyze the data reported in this paper is available from the lead contact upon request.

### EXPERIMENTAL MODEL AND SUBJECT DETAILS

#### Animal Model

All animal experiments were approved by the Institutional Animal Care and Use Committee and were conducted in compliance with the relevant ethical regulations. Mice were maintained in social cages on a 12-hour light/dark cycle with free access to food and water; animals were monitored daily for food and water intake. Wild-type CD1 mice were used to isolate primary cell cultures on postnatal day 3 (P03). Animals of both sexes were used in the analyses.

#### Human PSC lines and cell culture

Human PSCs were maintained on Geltrex-coated plates in Stemflex medium and a 5% CO2 environment at 37°C. Cells were passaged using Accutase in Stemflex supplemented with Chroman1 (2 μM), which was removed from the media the following day.

To generate genetically modified PGP1 and WTC11 iPSCs with the *ADNP* p.Tyr719* variant, we altered the nucleotide sequence NM_015339.4.2157C to G. Additionally, we inserted hemagglutinin (HA) tags immediately after Tyr719 to detect the resulting truncated proteins. In parallel, as a control, we inserted HA tags at the 3’ end of the endogenous ADNP gene in the PGP1 line. sgRNAs targeting proximal regions were designed using the CHOPCHOP online tool (https://chopchop.cbu.uib.no/) and cloned into the lentiCRISPRv1 plasmid (Addgene Plasmid #49535) using the Golden Gate Assembly protocol. The gene-targeting template plasmid used to generate the HA-tagged *ADNP* p.Tyr719* allele includes a puromycin-resistance cassette flanked by LoxP sites for Cre-recombinase mediated deletion, positioned after the HA tag sequence, and the 5′ and 3′ DNA homology arms. A similar gene-targeting template was used to generate the HA-tagged *ADNP* allele. The day before transfection, human iPSCs were dissociated into single cells with Accutase and plated at 250,000 cells per well in six-well plates containing StemFlex medium with Chroman1. For transfection, the template vector (1 μg) and the CRISPRv1-gRNA (0.5 μg) were incubated with Lipofectamine Stem Transfection Reagent (10 μl) and OptiMEM (250 μl, Invitrogen) for 10 minutes, then added dropwise to the iPSCs. Selection with puromycin (1μg/ml) in StemFlex medium started 24-48 hours after transfection and continued for approximately 10 days until stable colonies formed. Individual colonies were picked with a P200 pipette tip under a microscope. Heterozygous and homozygous ADNP p.Tyr719-HA* iPSC clones, as well as heterozygous ADNP-HA iPSC clones, were confirmed by PCR. The puromycin-resistance cassette was removed using Cre-recombinase.

To generate iPSCs with stable inducible ASCL1-DLX2 integration, we took advantage of a bi-directional tetracycline-inducible promoter via the PiggyBac transposon system, as previously described ^105^. The iPSCs were transfected with the PB-TO-ASCL1-DLX2 plasmid and the EF1a-transposase vector at a 2:1 ratio using Lipofectamine™ Stem Transfection Reagent. Selection with puromycin started 24-48 hours post-transfection and continued for 2 weeks. Successfully modified human iPSCs harboring stable TO-ASCL1-DLX2 integration were either cryopreserved for future applications or directly differentiated into neuronal cells. PB-TO-ASCL1-DLX2 (Addgene plasmid # 182307) and EF1a-transposase plasmid were gifts from iPSC Neurodegenerative Disease Initiative (iNDI) & Michael Ward.

Research on human-derived samples was conducted according to protocols approved by the institutional review boards of the Icahn School of Medicine at Mount Sinai.

#### Lentivirus production

Infectious lentiviral particles were produced in HEK293T cells using the third-generation lentiviral packaging plasmids: pMD2.G (Addgene plasmid # 12259), pRSV-rev (Addgene plasmid #: 12253), and pMDLg/pRRE (Addgene plasmid #: 12251) ^106^. Lentiviruses were produced in HEK293T cells using polyethylenimine (PEI), following established protocols ^107^. The lentiviral supernatant was centrifuged at 1,000g for 5 minutes to remove cellular debris. Lentiviral particles were then ultra-centrifuged, resuspended overnight with gentle shaking in DMEM, aliquoted, and stored at −80°C. Only virus preparations with >90% infection efficiency were used for experiments, as determined by GFP expression or antibiotic resistance.

#### Small molecule-mediated GABAergic neuron generation

Human PSCs were differentiated into cortical inhibitory neurons using an established protocol based on dual SMAD inhibition, WNT inhibition, and SHH activation,^42,43^ with the following modifications. NKX2.1(GFP/w) cells were dissociated into single cells using Accutase and plated at 430,000 cells/cm² onto Geltrex-coated wells in KSR media (415 mL Knockout DMEM, 75 mL knockout serum replacement, 500 µL 2-mercaptoethanol, 5 mL NEAA, and 5 mL GlutaMAX) supplemented with 10 µM Y-27632. From days 1 to 10, the cells underwent daily complete medium changes with LDN193189 (100 nM), SB431542 (10 µM), and XAV939 (2 µM). During days 11 to 18, N2 medium supplemented with 1 µM Purmorphamine and 5 µM recombinant SHH was used to induce ventral identity. On day 32, NKX2.1-GFP cells were dissociated, collected via FACS, and replated with mouse glial cells. The cells were maintained in Neurobasal A media supplemented with 1×B27, 1×Glutamax, 1% fetal bovine serum, and 2-4 µM AraC and harvested on day 67 for further analyses.

#### TF-mediated induced GABAergic neuron generation

We used our established protocol to generate GABAergic neurons through lentiviral-mediated overexpression of ASCL1 and DLX2, as previously described ^27^. Briefly, PSCs were transduced with FUW-TetO-Ascl1-T2A-puromycin, FUW-TetO-Dlx2-IRES-hygromycin, and FUW-rtTA lentiviruses. Twenty-four hours later, the medium was replaced by N2 media (1’N2 supplement, 1’NEAA in DMEM-F12 media) supplemented with 2 µg/mL doxycycline to induce transgene expression. The transduced cells were selected for three days by applying hygromycin (200 μg/mL) and puromycin (1 μg/mL). Subsequently, the media was replaced with N2 media containing 4 µM AraC for an additional three days. On day 8, GABAergic neurons were co-cultured with mouse glial cells at a density of 104,000 cells/cm^2^ on Geltrex-coated plates. Two weeks post-transgene induction, doxycycline was withdrawn, and the culture was maintained in Neurobasal A media supplemented with 1’B27, 1’Glutamax, and 1% fetal bovine serum. Mature neurons were used for various assays on day 35 after co-culture.

To generate GABAergic neurons from PSC lines with stable inducible ASCL1-DLX2 integration, PSCs were dissociated using Accutase and plated at a density of 100,000 cells/cm^2^ on GelTrex-coated plates in StemFlex Medium supplemented with Chroman1. The following day, the medium was replaced with N2 media containing 2 µg/mL Doxycycline to induce the expression of ASCL1 and DLX2. After three days, 4 µM AraC was added to the N2 media. On day 8, the neurons were replated with mouse glial cells. All subsequent procedures followed the previously described protocol for generating GABAergic neurons using lentiviral vectors.

#### Patterned induced GABAergic neuron generation

PSC lines with stable inducible ASCL1-DLX2 integration were dissociated into single cells using Accutase and plated at 430,000 cells/cm² onto Geltrex-coated wells in KSR media supplemented with 2 μM Chroman1. From days 1 to 10, the cells underwent daily full medium changes with LDN193189 (100 nM), SB431542 (10 μM), and XAV939 (2 μM). Starting on day 11, N2 media supplemented with 2 µg/mL doxycycline and 10 µM DAPT (Tocris Bioscience, 2634) was used to induce ASCL1 and DLX2 expression and promote synchronized neural differentiation. Doxycycline was discontinued either 3 or 7 days later, and DAPT was removed after 7 days. Neurons were then dissociated and replated with mouse glial cells. Subsequent steps followed the established protocols used for other GABAergic neuron differentiation methods.

### METHOD DETAILS

#### Primary mouse astrocytes

Mouse glial cultures were generated from cortical hemispheres at postnatal day 3 (P03). The cortices were incubated in 3 mL of 20 Units/mL Papain, 0.5 μM EDTA, and 1 μM CaCl2 in HBSS for 15 mins. After incubation, the tissues were manually dissociated by forceful trituration. The resulting cells were grown at 37°C with 5% CO_2_ in DMEM media containing 1×NEAA, 1× sodium pyruvate, 10% Cytiva HyClone™ Cosmic Calf™ Serum (CCS), and 0.008% β-mercaptoethanol.

#### Transplantation and Brain Slice Preparation

Concentrated cell suspensions (∼1 × 10^5^ cells/μL) were loaded into beveled glass micropipettes (Drummond Scientific). Each recipient received two injections of 0.5 μL into the forebrain at coordinates 2.0 mm posterior, 1.0 mm lateral, and 1.5 mm ventral to the skin surface, targeting the periventricular region. Stereotaxic coordinates were zeroed anteriorly at the inner corner of the eye, laterally at the midpoint between the eyes, and vertically at the skin surface. Following injection, pups were placed on a heating pad until fully warmed and active, then returned to their mothers. Animals were maintained with the dam until weaning at postnatal day 21 (P21).

Acute brain slices were prepared from approximately 6-month-old mice. Animals were deeply anesthetized with Euthasol and decapitated. Brains were rapidly removed and immersed in ∼0 °C aerated (95% O_2_ and 5% CO_2_) sucrose-based slicing solution containing (in mM): 213 sucrose, 2.5 KCl, 2 MgSO_4_, 2 CaCl_2_, 26 NaHCO_3_, 1.25 NaH_2_PO_4_, and 25 dextrose (315 mOsm, pH 7.4). In this slicing solution, coronal slices with a thickness of 300 mm were cut using a vibratome (Leica VT1200S). Slices were cut from both female and male mouse brain and then incubated in aerated artificial cerebrospinal fluid (ACSF) containing (in mM): NaCl 126, KCl 2.5, NaH_2_PO_4_ 1.25, NaHCO_3_ 26, dextrose 25, MgSO_4_ 2 and CaCl_2_ 2 (315 mOsm, pH 7.4) at ∼34°C for 50 min and then maintained at room temperature until recording.

#### Electrophysiological Recordings from Brain Slices

Slices were transferred to a recording chamber and perfused with oxygenated ACSF at 25°C. Human interneurons located in subcortical regions were visualized using an infrared differential interference contrast (IR-DIC) microscope (BX51WI, Olympus) equipped with a 40× water-immersion objective. Images were captured with a CMOS camera (C11440-36U, Hamamatsu) and displayed in real time. GFP-expressing human neurons were identified using 470-nm LED excitation. To assess intrinsic membrane properties, whole-cell current-clamp recordings were performed using 500-ms current steps of increasing amplitude. Patch pipettes (_∼_5 MΩ) were filled with a K-gluconate–based internal solution containing (in mM): 140 K-Gluconate, 3 KCl, 2 MgCl_2_, 0.2 EGTA, 10 HEPES and 2 Na_2_ATP (287 mOsm, pH 7.22), supplemented with 0.2% biocytin for post hoc morphological analysis. Voltage signals were filtered at 10 kHz and sampled at 40 kHz, while synaptic currents were low-pass filtered at 3 kHz and sampled at 40 kHz using a Multiclamp 700B amplifier and Clampex 11.2 software (Molecular Devices).

#### Whole-cell recordings from cultured neurons

Functional properties of iNs were assessed using whole-cell patch-clamp electrophysiology as previously described.^27,30^ Spontaneous inhibitory postsynaptic currents (sIPSCs) were recorded at a holding potential of −70 mV in the presence of 20 μM CNQX. Miniature inhibitory postsynaptic currents (mIPSCs) were recorded at −70 mV in the presence of 1 μM tetrodotoxin (TTX) and 20 μM CNQX. Patch pipettes were filled with a Cs-based internal solution containing (in mM): 40 CsCl, 3.5 KCl, 10 HEPES, 0.05 EGTA, 90 K-gluconate, 1.8 NaCl, 1.7 MgCl₂, 2 Mg-ATP, 0.4 Na-GTP, and 10 phosphocreatine. The external recording solution contained (in mM): 140 NaCl, 5 KCl, 10 HEPES, 2 CaCl_₂_, 2 MgCl_₂_, and 10 glucose (pH 7.4). All recordings were performed at room temperature.

#### Immunofluorescence and imaging

Cultured cells were fixed with 4% paraformaldehyde in D-PBS at room temperature for 5-10 min. Cells were rinsed 3 times with D-PBS and subsequently permeabilized for 5 mins at room temperature with 0.2% Triton X-100 in D-PBS. After incubation in blocking buffer (4% bovine serum albumin (BSA) and 1% Cosmic Calf™ Serum in D-PBS) for 3 hours, primary antibodies were added for incubation overnight at 4^°^C. After 3 rinses, cells were incubated with secondary antibodies in the blocking buffer for 3 h at room temperature. Images were acquired using the EVOS M5000 imaging system (Life Technologies).

Images of injected human neurons in mouse brain sections were acquired at the Microscopy and Advanced Bioimaging CoRE at the Icahn School of Medicine at Mount Sinai. Imaging was performed using ZEN Blue software (v3.7.4) on a Zeiss LSM980 confocal microscope equipped with a 32-channel Airyscan 2 GaAsP-PMT detector, operated in super-resolution (SR) mode with Multiplex 4Y. Images were collected using a 100×/1.46 NA α-Plan-Apochromat oil-immersion objective with a minimum optical zoom of 1.7. Frame size was set to “optimal” to ensure Nyquist sampling for the selected field of view. Images were acquired at 16-bit depth, with scan speed set to “maximum” and a frame average of 1. Excitation was provided by an argon 488-nm laser for GFP and a 561-nm diode laser for Alexa Fluor 561. Emission was collected using a main beam splitter (MBS) 488/561, a secondary beam splitter (SBS) SP615, and dual emission filters (BP 420–480 and BP 495–550 for GFP; BP 570–620 and LP645 for Alexa Fluor 561). For z-stack acquisition, the z-step size was set to half of the axial resolution (0.4 µm per step) to ensure adequate sampling. For each experimental cohort, positive control samples were used to establish excitation and detection parameters, including laser power and detector gain. Settings were optimized to maximize signal-to-noise ratio while using approximately 50% of the detector’s dynamic range and avoiding saturation. Detector gain was maintained below 850 V to ensure linear response, and laser power rarely exceeded 1%. Once established, these parameters were held constant for all samples within each cohort. Image processing was performed using the ZEN Airyscan processing module.

#### Immunoblotting

Cells were washed three times with cold PBS, harvested using CELLTREAT cell lifters, and lysed with the RIPA Lysis and Extraction Buffer (supplemented with 1X Halt Protease and Phosphatase Inhibitor Cocktail and 5 mM EDTA). Cells were incubated on ice for 15 min with occasional vortexing, sonicated twice (1s pulse followed by 1 min on ice), incubated on ice for another 15 min with occasional vortexing, and then centrifuged at 21000 xg for 20 min with the supernatant collected for quantifications using the BCA assay. Samples were denatured with 1X NuPAGE LDS loading buffer (supplemented with 50 mM DTT) and incubated at 70 °C for 10 min.

Approximately 25 µg of total proteins were loaded onto each well of the 4-12% Bis-Tris gel and run at 170 V for 30 min in 1X MOPS Buffer. Samples were then transferred onto PVDF membrane at 20 V for 1 h at 4 °C in cold 1X Transfer Buffer (25 mM Tris, 200 mM Glycine, 15% MeOH in ddH2O), blocked with either the LICOR Blocking Buffer or 5% milk in TBST for 1h at RT, then incubated overnight at 4°C on the rocking shaker with the addition of primary antibodies diluted in the corresponding blocking buffer supplemented with 0.1% Tween-20. Membranes were washed four times with TBST (5 min at RT on the shaker each time), incubated at RT for 1h with secondary antibodies (diluted in the corresponding blocking buffer and supplemented with 0.1% Tween-20 and 0.1% SDS), washed for another 4 times with TBST (for HRP antibodies, treated with the ECL buffer at A:B = 1:1 in dark for 5 min at RT before imaging), and imaged using the Bio-Rad ChemiDoc MP Imaging System.

#### 10X single-cell RNA-seq library preparation and sequencing

We used a Papain solution to create single-cell suspensions based on a published method.^108^ Neurons were detached from the plate using Accutase supplemented with 0.4% DNase (Worthington), collected and further dissociated with a Papain solution. This solution was prepared by adding 328 µL of Papain to each 20 mL Enzyme Stock Solution containing 1× EBSS, 0.46% D(+) Glucose, 26 mM NaHCO3, and 0.5 mM EDTA. Additionally, 0.004 g of L-cysteine was added before use. The solution was activated for 15 minutes at 37°C, followed by an additional 1.5-hour incubation at 37°C. Prepare the Inhibitor Stock Solution (0.46% D(+) Glucose and 25 mM NaHCO3 in1× EBSS), Low Ovo (10X) solution (3 g BSA (Sigma A8806), and 3 g Trypsin inhibitor (Worthington LS003086) in 200 mL D-PBS, adjusting the pH to 7.4) and High Ovo (5X) solution (6 g BSA and 6 g Trypsin inhibitor in 200 mL D-PBS, pH 7.4). The single-cell suspension was washed four times, alternating between low Ovo solution (9 mL inhibitor stock solution and 1 mL Low Ovo (10X)) and high Ovo solution (4 mL inhibitor stock solution and 1 mL High Ovo (5X)). After washing, the cells were centrifuged at 350 g for 5 minutes, and the resulting cell pellet was resuspended in cold 0.02% BSA. Single-cell RNA sequencing library preparation was performed using the Chromium Single Cell 3’ Reagent V3 kit, following the manufacturer’s specifications. Barcoded libraries were sequenced on the Illumina NovaSeq platform with 150 bp paired-end reads, achieving an average depth of approximately 30,000 reads per cell.

#### SHARE-seq library preparation and sequencing

SHARE-seq libraries were prepared as previously described^67^. Briefly, cells were fixed using 0.2% formaldehyde and permeabilized. For joint measurements of single-cell chromatin accessibility and expression (scATAC- and scRNA-seq), cells were first transposed by Tn5 transposase to mark regions of open chromatin. The mRNA was reverse transcribed using a poly(T) primer containing a unique molecular identifier (UMI) and a biotin tag. Permeabilized cells were distributed in a 96-well plate to hybridize well-specific barcoded oligonucleotides to transposed chromatin fragments and poly(T) cDNA. Hybridization was repeated three times to expand the barcoding space and ligate cell barcodes to cDNA and chromatin fragments. Reverse crosslinking was performed to release barcoded molecules. cDNA was separated from chromatin using streptavidin beads, and each library was prepared separately for sequencing. Libraries were sequenced on the Illumina NovaSeq platform (read 1: 50 cycles, read 2: 50 cycles, index 1: 99 cycles, index 2: 8 cycles).

### QUANTIFICATION AND STATISTICAL ANALYSIS

#### 10X Single-cell RNA-sequencing data processing and analysis

All single-cell RNA sequencing (scRNA-seq) samples were processed individually. Sequencing reads were aligned to a combined GRCh38 human and mm10 mouse reference transcriptomes (GRCh38_and_mm10-2020-A) using Cellranger (v7.0) with default parameters to generate gene-by-cell count matrices. Downstream analyses were performed in R using Seurat (v5.0.1). The data was filtered to retain only genes annotated to the human genome, and features were retained if detected in at least 3 cells and with a minimum of 200 features per cell. High-quality cells were selected based on having 500 to 7,000 detected genes and a maximum of 20% of reads mapping to mitochondrial genes. For 7D-patterned neurons and ADNP neuron samples, doublets were removed using DoubletFinder (v2.0) with default parameters. After preprocessing, samples were merged and normalized using natural-log transformation, and counts were scaled by a factor of 10,000. Principal component analysis (PCA) was conducted on the 2,000 most variable features (FindVariableFeatures()). The first 20 principal components (k=20) were used to construct a K-nearest neighbor (KNN) graph, while the first 30 principal components were used for ADNP samples. Cells were clustered using the Louvain algorithm (FindClusters()) and visualized using UMAP embeddings. Clusters were manually annotated with cell types based on a curated list of markers (Table S2). When necessary, samples were integrated using the FastMNN algorithm (IntegrateLayers()). After merging or integration, genes related to the cell cycle and ribosomal function were regressed out. Differential expression analysis between clusters or samples was performed using the Wilcoxon rank sum test using only genes expressed by a minimum of 3-5% of cells. Method/cluster specificity genes were identified as previously described ^109^. Briefly, genes with non-unique names or not expressed in any cells were filtered out. Gene expression was scaled to 1 million unique molecular identifiers per method or cluster. Gene expression specificity was calculated by dividing the expression of each gene in each method/cluster by the total expression of all genes in that method/cluster, resulting in values from 0 to 1, with values closer to 1 indicating high specificity. The top 10% of the most highly specific genes per method or cluster were used for gene expression analyses. Pseudotime trajectory analysis was performed using Slingshot (v2.7.0) or Monocle3 (v.1.3.7) software with default parameters. Graphtest() in Monocle3 was used to determine genes that change with pseudotime.

To assess the similarity of PSC-derived GABAergic neurons to inhibitory neurons from different regions of the developing fetal brain, we utilized previously published scRNA-seq data from Bhaduri et al. ^53^ encompassing weeks 18, 19, 20, and 25 of gestation across various brain regions (neocortex, striatum, thalamus, hypothalamus, ganglionic eminence, midbrain, and cerebellum). Additionally, hypothalamic neuron data from Herb et al. ^54^ were included. All datasets were merged into a single Seurat object for comprehensive analysis. The combined dataset was first refined to focus on neurons expressing the inhibitory neuron makers GAD1/2 (log norm counts > 0). We employed Seurat’s FindTransferAnchors() and MapQuery() functions for cell mapping and comparison. The data normalization was carried out using SCTransform.

To compare cell states across differentiation approaches, publicly available scRNA-seq datasets were independently reprocessed using Seurat. Gene-by-cell count matrices were log-normalized and scaled, and the top 2,000 highly variable genes were selected for dimensionality reduction. Principal component analysis (PCA) was performed using 50 components, followed by UMAP embedding (min.dist = 0.3). To assign lineage and cell-state identities, each dataset was mapped onto a reference atlas of the developing human brain.^49^ The reference dataset was processed using the same pipeline, except that 20 principal components were used for UMAP construction. Anchor-based mapping was performed using FindTransferAnchors() (reduction = “rpca”, k.anchor = 20), followed by cell-type annotation with TransferData() (dims = 1:20, k.weight = 10). This strategy enabled consistent assignment of interneuron lineage and regional identities across differentiation methods.

#### SHARE-seq data processing and analysis

SHARE-seq libraries were aligned using a published pipeline (https://github.com/masai1116/SHARE-seq-alignmentV2). Briefly, sequencing reads were trimmed and aligned to the hg38 genome using Bowtie2. Reads were demultiplexed using four sets of 8-bp barcodes in the index reads, allowing one mismatch per barcode. Reads mapping to the mitochondria and chrY were discarded. Duplicates were removed based on the read alignment position and cell barcodes. Open chromatin region peaks were called on merged samples using MACS2. Peaks overlapping with ENCODE blacklisted regions (https://sites.google.com/site/anshulkundaje/projects/blacklists) were filtered out. Peak summits were extended by 250bp on each side and defined as accessible regions. The fragment counts in peaks and TF scores were calculated using chromVAR. SHARE-RNA-seq reads were aligned to the hg38 genome using STAR. Reads were demultiplexed as described above for SHARE-ATAC-seq. Aligned reads were annotated to both exons and introns using feature counts. PCR duplicates were removed based on UMIs and read alignment position. Cells that expressed >1,000 genes and <4% mitochondrial reads were retained. Seurat was used to preprocess the count matrix and cluster cells for scRNA-seq libraries as described above. For pseudotime trajectory analysis, hPSCs, NPC-like cells, and stressed cells were removed, and Slingshot was utilized with default parameters.

#### Inferring regulatory networks

To infer transcription factor (TF) regulon networks within our single-cell RNA-seq dataset, we utilized the SCENIC (Single-Cell Regulatory Network Inference and Clustering) ^61^ protocol (v1.1.2), which employs the vsn-pipeline version orchestrated via Nextflow with the pyscenic (v0.12.1) container. First, Seurat objects were converted into loom files using the SCopeLoomR package (v0.13.0). Subsequently, the pyscenic module was executed 20 times using its default settings. Only regulons identified in at least 15 out of the 20 runs were included for further analysis.

#### Gene Ontology, pathway enrichment and disease-vulnerable cell type analysis

We used Enrichr ^110^ or ClusterProfiler (with universe set to all expressed genes in dataset to identify the cellular functions and biological processes of differentially expressed genes. To identify the cellular functions of differential peaks, we annotated the peaks to their nearest genes, followed by Enrichr analysis.

To determine the enrichment of genes implicated in autism spectrum disorder (ASD), we used the Simons Foundation Autism Research Initiative (SFARI) database to compile a list of 1,065 ASD-associated genes scored 0-2 or categorized as syndromic. Fisher’s exact test was performed to determine the statistical significance of the top 10% of differentially expressed genes specific to PSC-iGABAs or patterned-iGABAs.

For a more comprehensive analysis of all neural-related diseases, we used the ToppGene Suite and selected the DisGeNET Curated database,^57^ which includes a total of 11,181 annotations. DisGeNET integrates information on gene-disease associations from repositories, GWAS catalogs, animal models, and scientific literature. To control for multiple testing errors, a threshold of 0.05 was applied. We used a random sampling size of 5,000, considering only coding genes to ensure the results were not biased by the size of the gene list, and required a minimum feature count of at least two genes in our DEG list. Only disease associations with an FDR-adjusted p-value ≤ 0.05 (Benjamini-Hochberg for multiple hypothesis testing) were considered significant.

To assess disease vulnerability across neurons generated using our differentiation approaches, we integrated our scRNA-seq data with publicly available reference datasets from postmortem human brain tissue. Specifically, we used datasets from individuals with AD^59^ and ASD^60^, which define interneuron populations reported to exhibit selective vulnerability in each disease context. Using a reference-mapping framework, we projected cells from these datasets onto our induced GABAergic neuron atlas. We then quantified the fraction of cells generated by each differentiation strategy that mapped to disease-vulnerable cell types, enabling comparative assessment of how well each approach captures interneuron populations implicated in AD- and ASD-related pathology.

#### WGCNA analysis

A total of 8,516 genes with an average expression level above 0.15 were selected for WGCNA analysis.^111,112^ To construct the network, a soft thresholding is selected through scale independence: power = 4 for clusters 0 and 2 and power = 3 for cluster 1. Gene clustering and module assignments were visualized using a dendrogram, highlighting the hierarchical relationships between gene modules.

#### Statistical analyses

Unless otherwise indicated, all data presented are the average of at least two biological replicates from each of at least two independent experiments. Statistical analysis was matched to the data structure as noted above in the Methods Details section for single-cell RNA-seq and SHARE-seq experiments. Statistical analyses were performed in RStudio (v4.3.1; RStudio team, 2020) or GraphPad Prism 9. See figure legends for details on specific statistical tests run for each experiment. Statistical significance is represented by a star (*) and indicates a computed P-value < 0.05. Graphs and plots were generated using Graphpad Prism or RStudio. Figures were generated using Adobe Illustrator and Biolegend.

