## Supplementary Figures for "Expanding GABAergic Neuronal Diversity in PSC-Derived Disease Models"

**Figure S1. Generation of Immature Cortical GABAergic Neurons from Human PSCs Using A Directed Differentiation Protocol.**

*Related to Figure 1.*

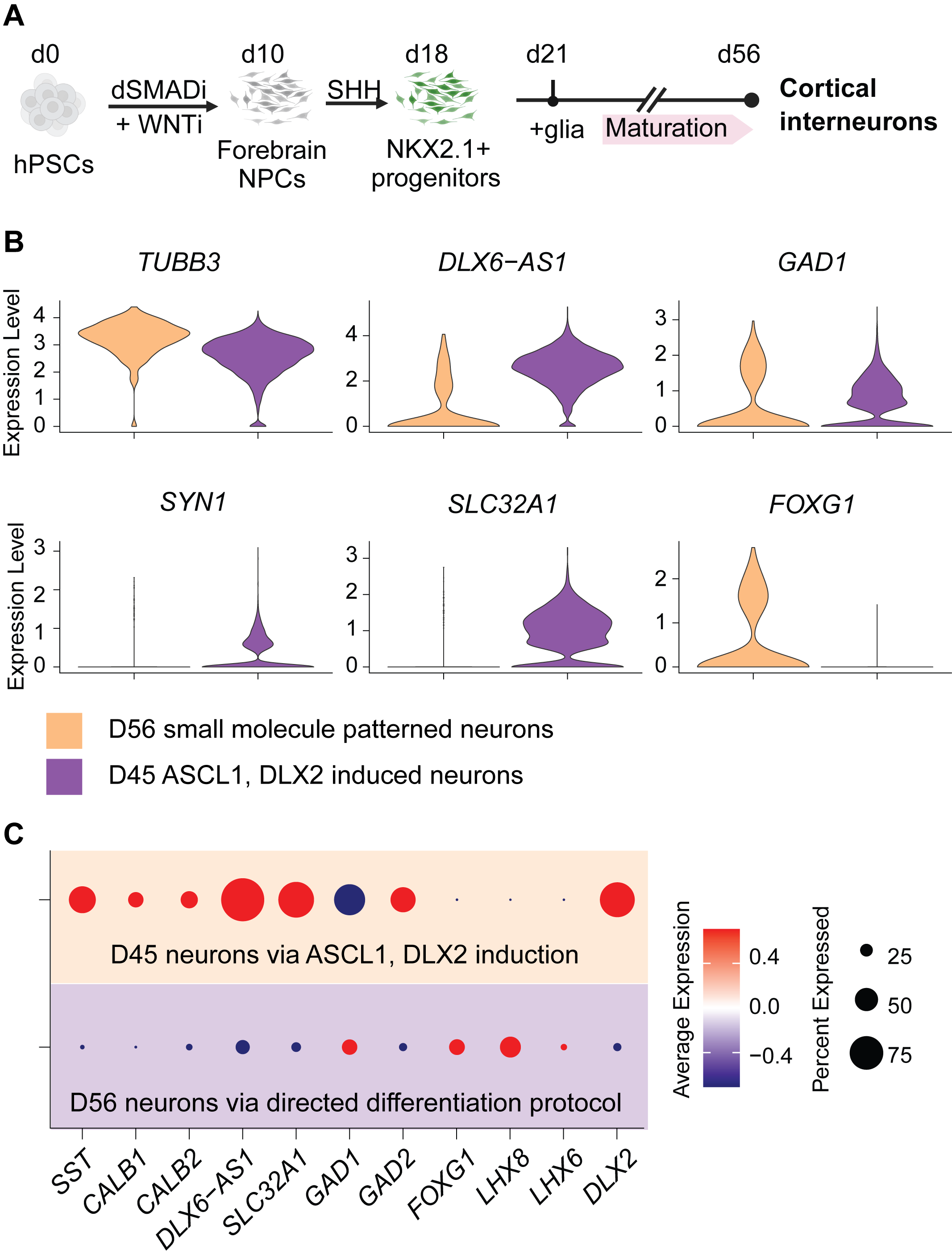

**Figure S1.** (**A**) Schematic overview and timeline of the small molecule method, followed by single-cell transcriptomics analysis. (**B-C**) Violin plots (**B**) and dot plots (**C**) show expression levels of neuronal identity and maturation-related marker genes in GABAergic neurons generated using either transcription factors (TFs), i.e., ASCL1 and DLX2, or small molecules.

**Figure S2. Induction of Neuronal Fate Using ASCL1/DLX2 and Patterning Factors.**

*Related to Figure 1.*

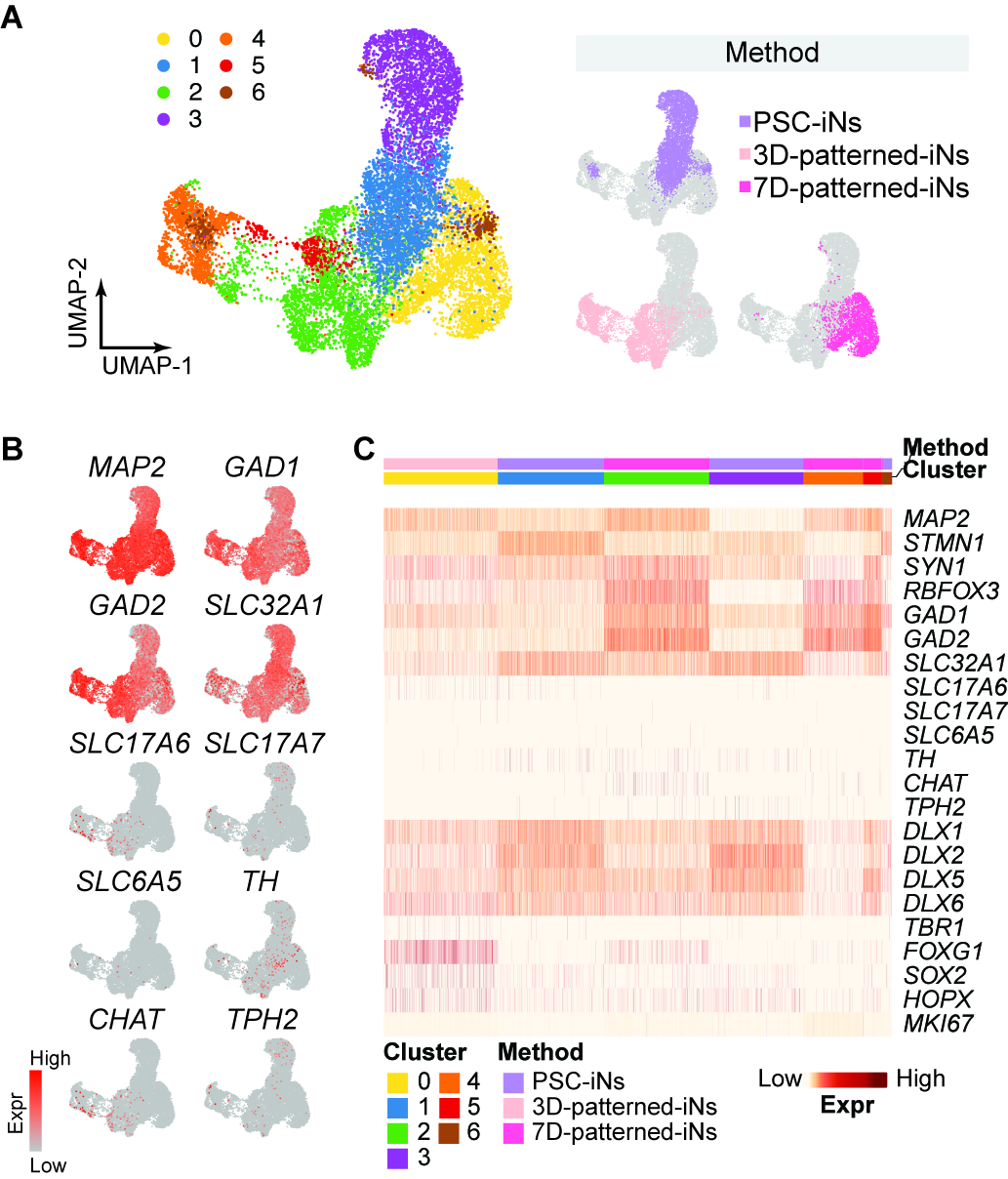

**Figure S2.** (**A**) UMAP embedding of 12,876 cells, color-coded by cluster identity (left panel) and differentiation method (right panel). (**B**) Feature plots for neuron subtype-specific markers. (**C**) Heatmap of cell type-specific marker gene expression across clusters. See also Table S1.

**Figure S3. Expression of Neuronal Subtype, Maturation, and Brain Regionality Marker Genes in GABAergic-Induced Neurons.**

*Related to Figure 1.*

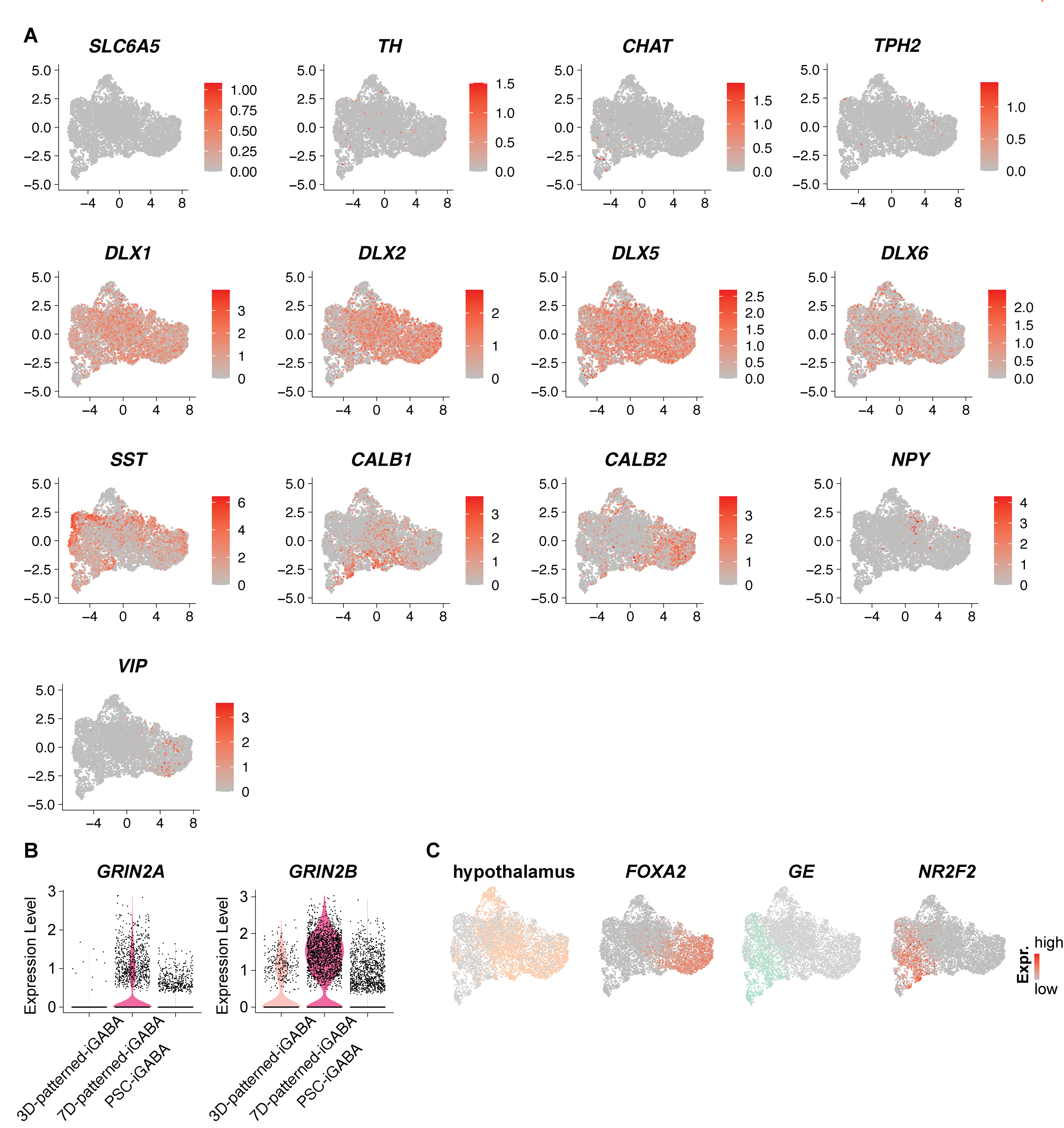

**Figure S3. (A)** Feature plots showing the expression of additional neuronal subtype marker genes in GABAergic-induced neurons. **(B)** Violin plots displaying the expression levels of NMDA receptor subunits in GABAergic neurons generated using different methods. **(C)** UMAP visualization depicting the respective brain regions and relevant TFs. See also Figure 1.

**Figure S4. Comparison of Interneuron Subtype Composition Across Differentiation Methods.**

*Related to Figure 1.*

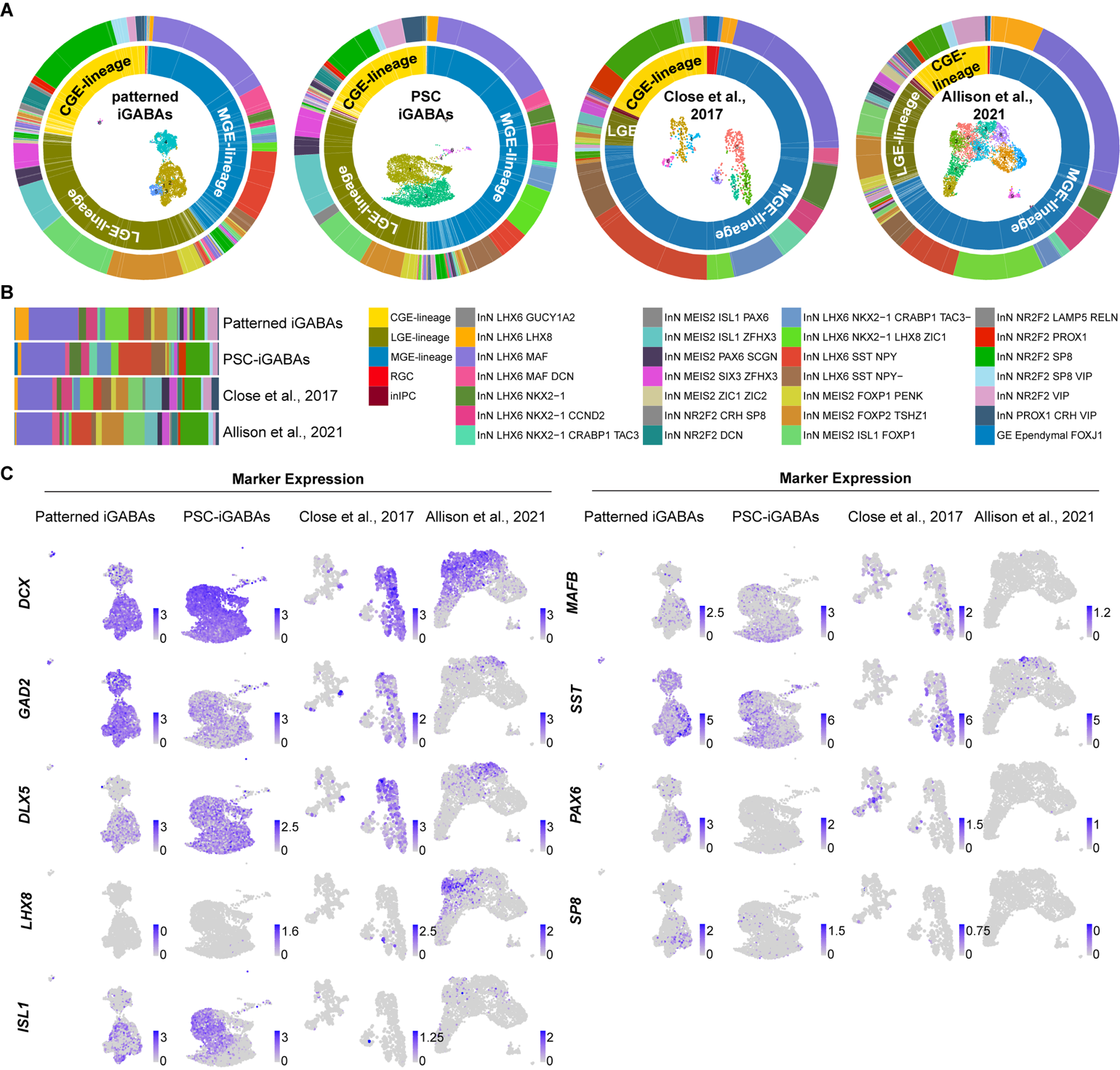

**Figure S4.** (**A**) Transcriptomic mapping of human PSC-derived interneurons generated using four protocols, patterned GABAergic-iNs (this study), GABAergic PSC-iNs (this study), Close et al. (2017), and the Close et al. (2017) and Alison et al. (2021) method, to reference human brain interneuron datasets. The outer rings indicate the relative abundance of mapped interneuron subtypes. (**B**) Proportional representation of major interneuron lineage groups across the different methods, highlighting their differing patterning tendencies. (**C**) Expression of key interneuron markers in each dataset, confirming molecular identities consistent with multiple interneuron origins. See also Figure 1.

**Figure S5. In Vivo Survival and Neuronal Differentiation of Transplanted Patterned-iNs.**

*Related to Figure 3*

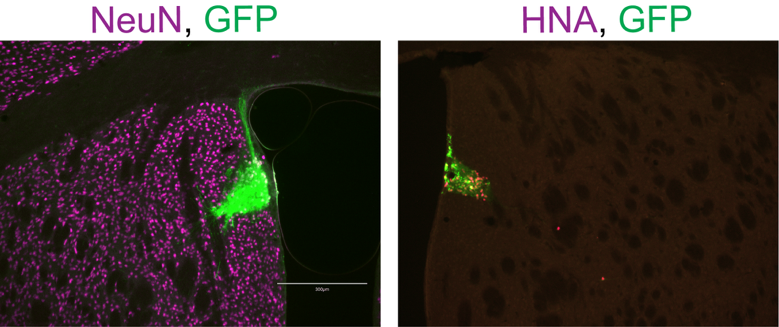

**Figure S5.** At 5 months post-transplantation, immunohistochemical analysis revealed that the majority of transplanted human cells, identified by human nuclear antigen (HNA) and GFP expression, differentiated into NeuN⁺ neurons. Most engrafted cells were detected in subcortical areas adjacent to the injection site, including the lateral septum and striatum. Scale bar: 300µm. These findings indicate long-term survival and stable neuronal differentiation of patterned-iNs following neonatal xenotransplantation. See also Figure 3.

**Figure S6. Disease Gene Enrichment in GABAergic PSC-iGABAs and patterned-iGABAs.**

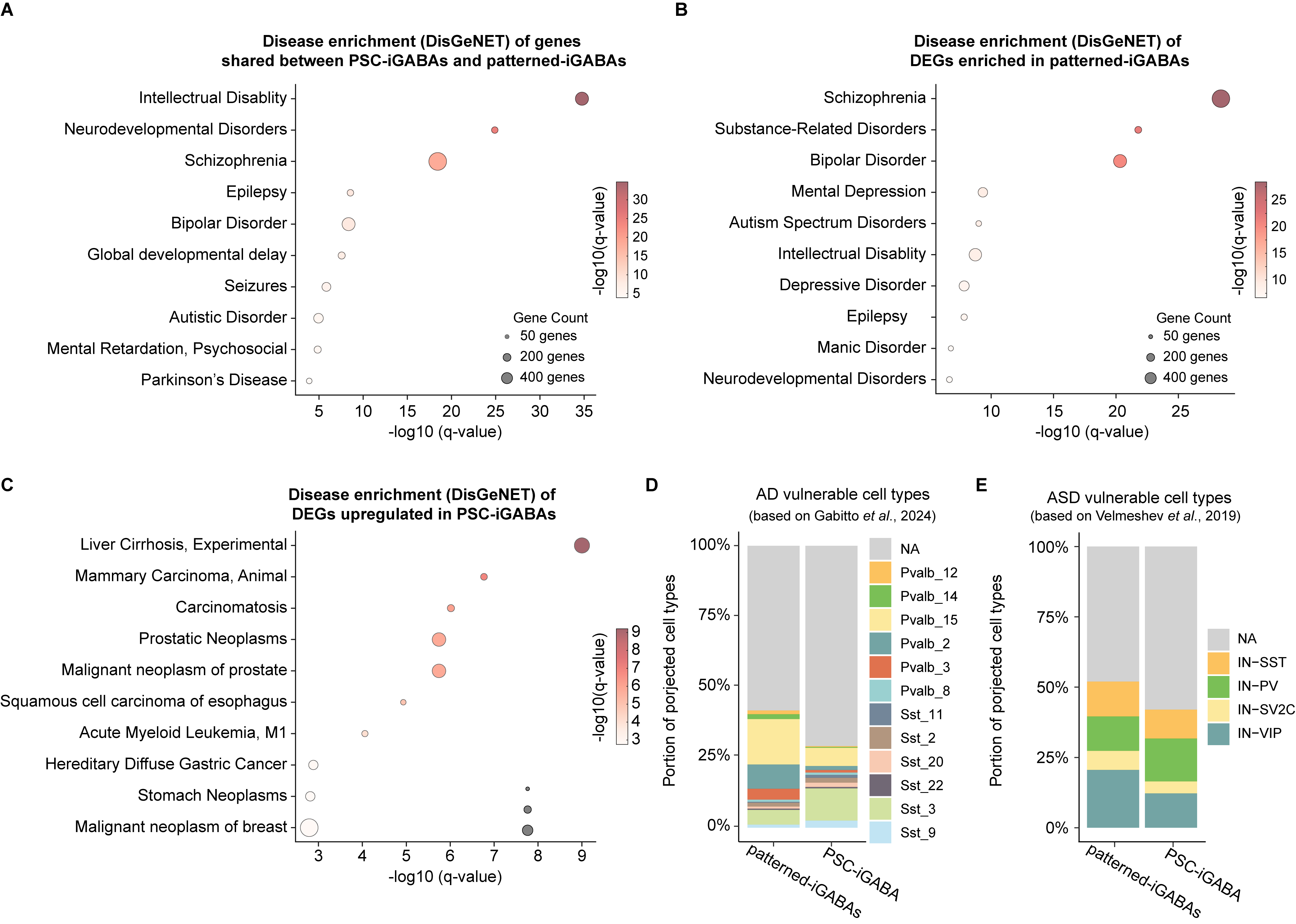

**Figure S6. (A–C)** Disease enrichment analysis of differentially expressed genes (DEGs) using the DisGeNET platform. **(A)** Diseases associated with DEGs shared between PSC-induced GABAergic neurons (PSC-iGABAs) and patterned induced GABAergic neurons (patterned -iGABAs). **(B)** Diseases enriched in DEGs specific to patterned-iGABAs. **(C)** Diseases enriched in DEGs specific to PSC-iGABAs. The x-axis represents the gene count for each disease, and dot size indicates the number of genes associated, while color intensity reflects –log10(p-value) enrichment significance. **(D, E)** Assessment of disease-vulnerable neuronal subtypes represented in the two differentiation methods. **(D)** Cell-type overlap analysis of GABAergic neuron subtypes generated by patterned-iNs and PSC-iNs with AD-vulnerable cell types (based on Mathys et al., 2019). **(E)** Similar analysis comparing subtype overlap ASD-vulnerable interneuron types (based on Velmeshev et al., 2019). These results demonstrate that both differentiation strategies can generate interneuron subtypes implicated in neurodevelopmental and neurodegenerative disorders, providing a foundation for disease modeling applications.

**Figure S7. Gene Regulatory Network (GRN) analysis centered on method-specific modulators.**

*Related to Figure 4.*

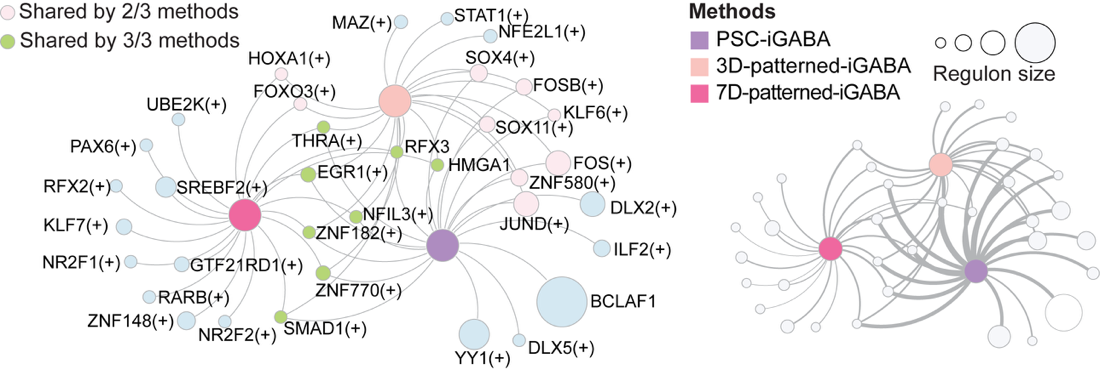

**Figure S7.** Left: Regulons are color-coded based on uniqueness, with node size proportional to regulon size, indicating the number of target genes regulated by each TF. Right: The same GRN with edge thickness represents the strength of the association between method modulators and regulons. See also Figure 4.

**Figure S8. Differential Expression of NPC Genes and Their Induction Along the Differentiation Trajectory of PSC-iNs and patterened-iNs.**

*Related to Figure 5.*

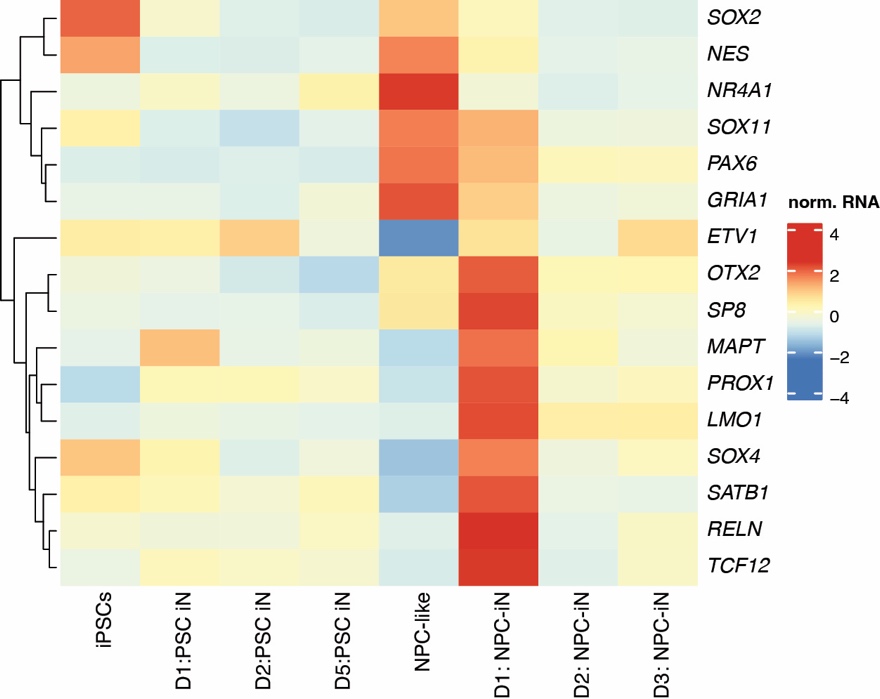

**Figure S8.** Heatmap showing gene expression across pseudobulk samples, aggregated from individual differentiation stages along the PSC-iNs and patterned iNs trajectory. See also Figure 5.

**Figure S9. Non-interneuron Lineage Genes Are Minimally Induced in GABAergic PSC-iNs and Patterned-iNs.**

*Related to Figure 5*

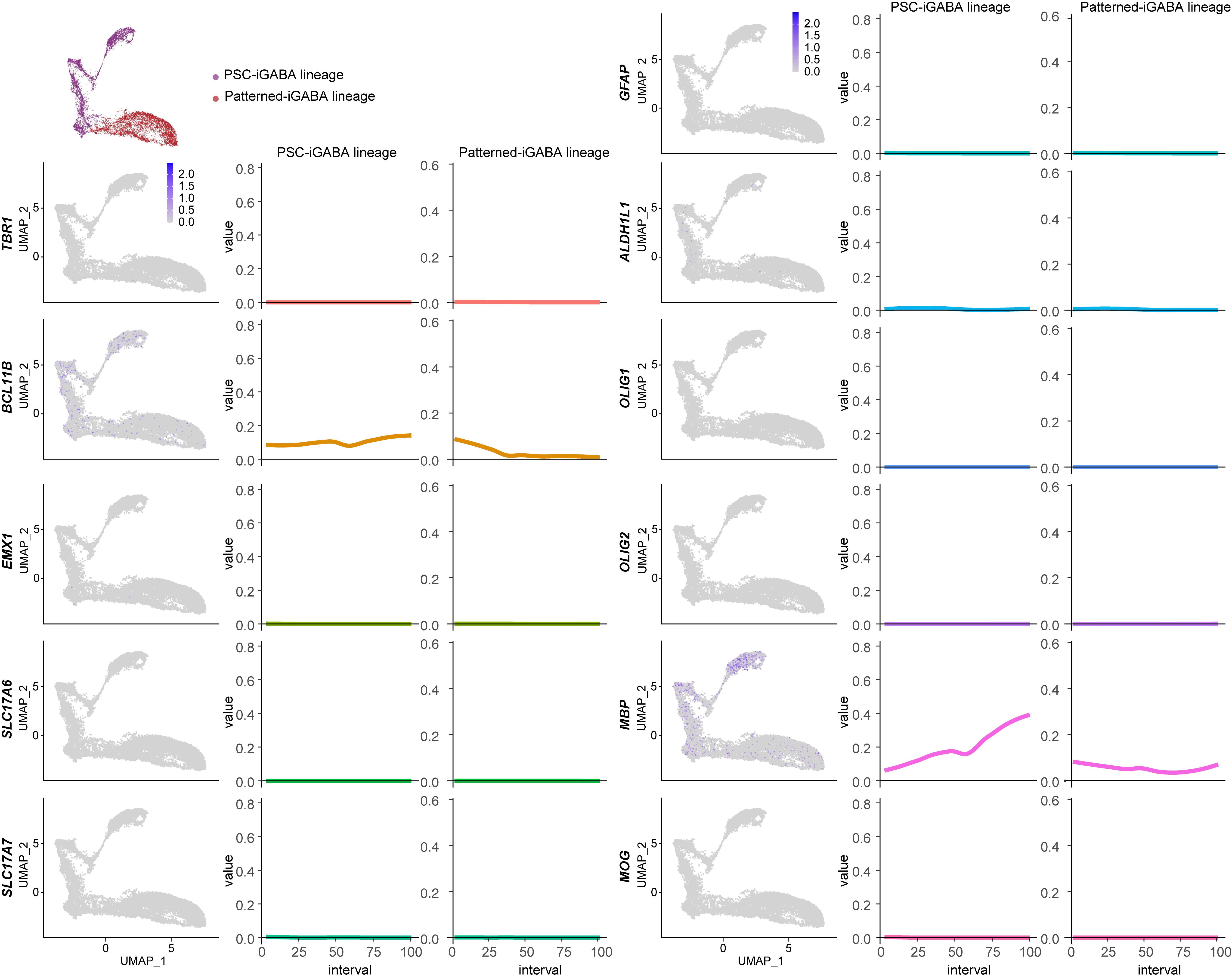

**Figure S9.** UMAP and pseudotime trajectory plots showing expression dynamics of non-GABAergic lineage markers in PSC-derived GABAergic neurons (PSC-iGABAs) and patterned induced GABAergic neurons (patterned-iGABAs). Expression of genes associated with alternative lineages, including *GFAP*, *ALDH1L1* (astrocyte), *TBR1*, *BCL11B*, *EMX1*, *SLC17A6*, *SLC17A7* (excitatory neurons), and *OLIG1*, *OLIG2*, *MBP*, *MOG* (oligodendrocyte), was analyzed across the differentiation trajectory. Expression of these non-interneuron markers remained low or undetectable across both lineages, indicating minimal contamination by alternative cell fates. Notably, MBP showed mild induction in the PSC-iGABA lineage but not in patterned-iGABAs, suggesting a slightly greater propensity for glial gene activation in PSC-derived cells. These data support the lineage specificity of both differentiation protocols, with rare off-target induction of non-interneuron identities. See also Figure 5.

**Figure S10. Generation of iPSC Lines with HA-Tagged Wild-Type ADNP or ADNP^p.Tyr719.*^**

*Related to Figure 6.*

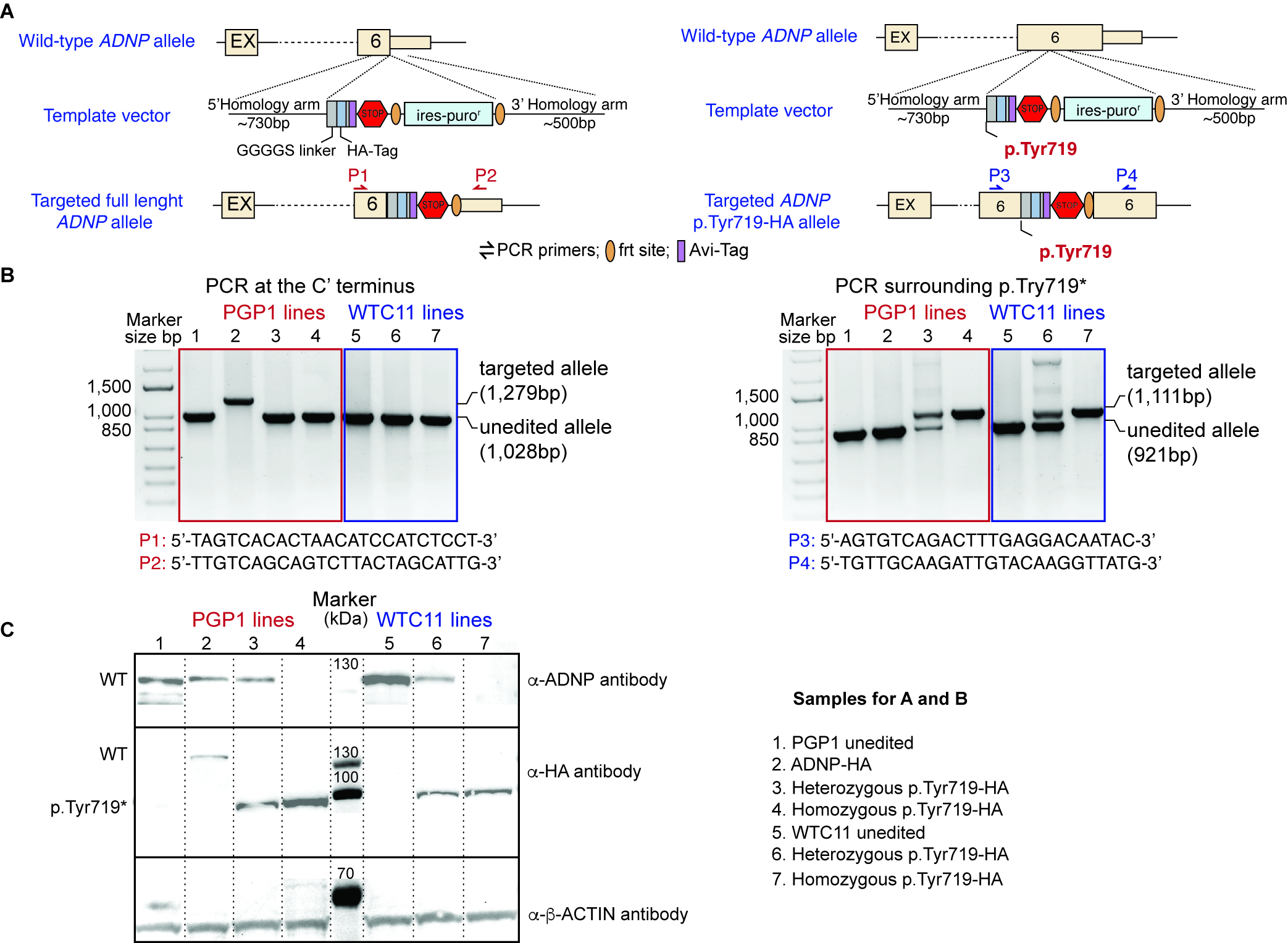

**Figure S10.** (**A**) Schematic representation of the generation of HA-tagged wild-type ADNP (left) and ADNP^p.Tyr719*^ (right) iPSC lines. The targeting vectors included two homology arms flanking a selection cassette containing an internal ribosomal entry site (IRES), a puromycin resistance gene (puro), and a polyadenylation signal. The selection cassette was flanked by two flippase recognition target (FRT) sites, and FlipE recombinase was expressed to remove the selection cassette. (**B**) Validation of homologous recombination by PCR. The HA-tagged wild-type allele (left) produced a band of approximately 1.2 kb, while the wild-type allele produced a band of approximately 1 kb. The HA-tagged ADNPp.Tyr719* allele produced a band of approximately 1.1 kb, and the unedited allele produced a band of approximately 900 bp. Primer sequences are provided. (**C**) Immunoblotting analysis confirmed the expression of full-length ADNP and ADNP^p.Tyr719*.^ β-ACTIN was used as the loading control. See also Figure 6.

**Figure S11.** ***ADNP^p.Tyr719*^* Mutation Disrupts Neurogenesis.**

*Relate to Figure 6.*

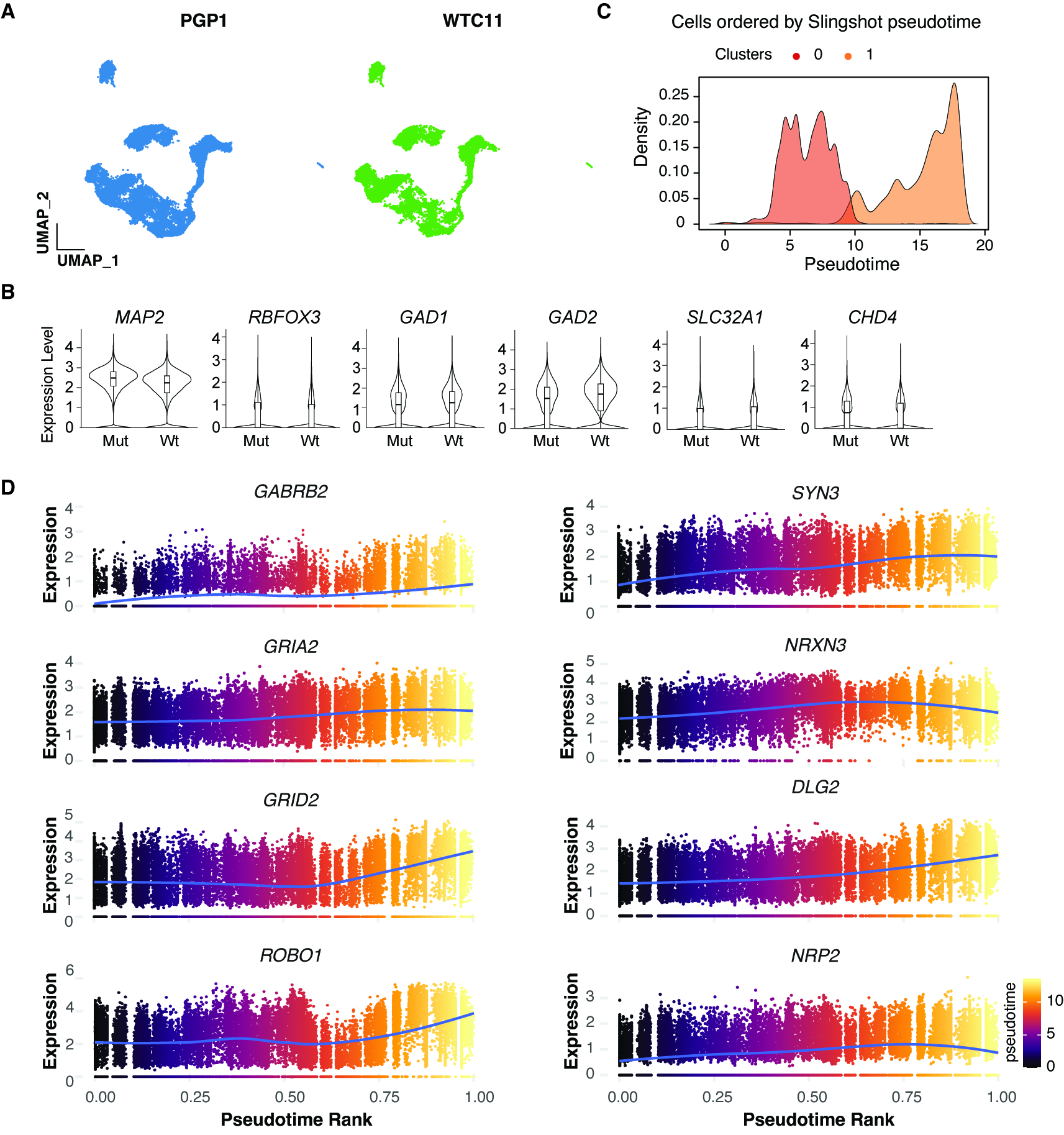

**Figure S11.** (**A**) Combined UMAP visualization of cells derived from different donor iPSCs. (**B**) Violin plots displaying the expression levels of pan-neuron and GABAergic neuron marker genes. (**C**) Inferred pseudotime analysis showing the progression from cluster 0 to cluster 1. The panel shows the density of each cluster across pseudotime. (**D**) Pseudotime expression plots show an increasing trend in genes related to neurite growth and synapse formation as cells progress along the pseudotime trajectory. Related to Figure 6.

**Figure S12. WGCNA Reveals Disruption of Gene Expression Networks Involved in Neuronal Development and Function in Mutant Cells.**

*Related to Figure 6.*

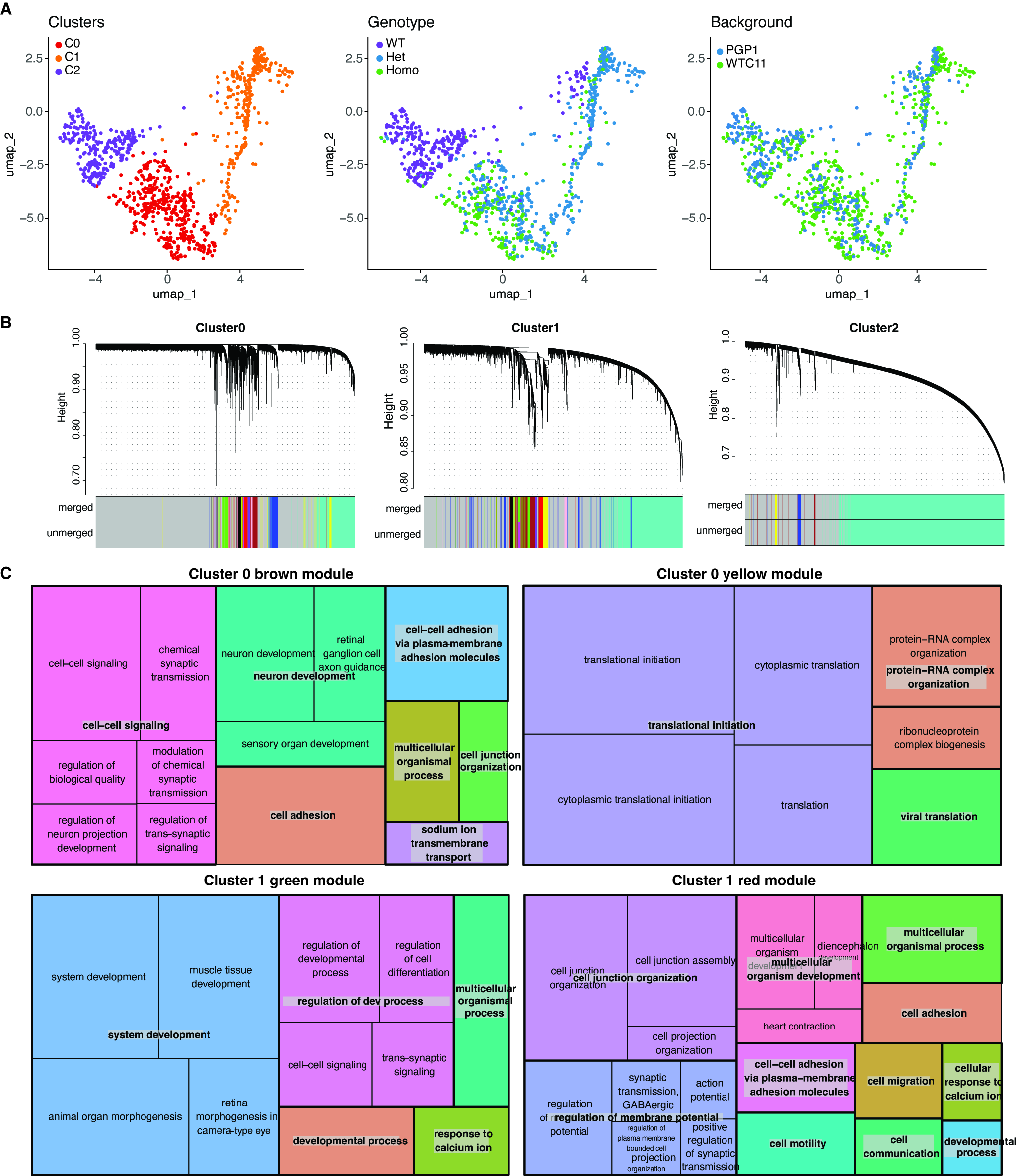

**Figure S12. (A)** Metacells maintain the cluster structure and accurately represent the genotypes and iPSC donor identities. **(B)** Gene expression modules identified within clusters 0, 1, and 2. **(C)** Treeplots displaying GO terms for biological processes associated with different modules in clusters consisting of mutant cells. See also Figure 6.
